# Pharmacological characterisation of allosteric modulators at human mGlu_5_

**DOI:** 10.1101/2024.12.05.627101

**Authors:** Muhammad Abdur Razzak, Kevin Tran, Roisin McCague, Kathy Sengmany, Jackson Kos, Monica Langiu, Bohan Li, Shane D Hellyer, Karen J. Gregory

## Abstract

The metabotropic glutamate receptor 5 (mGlu_5_) is a Class C G protein-coupled receptor, ubiquitously expressed throughout the CNS. With major roles in cognition, learning and memory, mGlu_5_ dysfunction is linked with numerous neurodegenerative and neuropsychiatric disorders, presenting a viable therapeutic target. Allosteric modulators bind topographically distinct sites from glutamate and other orthosteric agonists and enhance (positive allosteric modulators, PAMs), inhibit (negative allosteric modulators, NAMs) or do not effect (neutral allosteric ligands, NALs) mGlu_5_ function. While mGlu_5_ modulators have efficacy in *in vivo* rodent models of CNS disorders, none have made it to the clinic. We hypothesise this lack of translatability is arises from preclinical optimisation using on non-human pharmacological data, as functional studies are predominantly performed using rat mGlu_5_ and non-human brain neuronal cultures. Here we assess and quantify the impact of eleven chemically and pharmacologically diverse mGlu_5_ PAMs, NAMs and NALs on human mGlu_5_ activity using radioligand binding, iCa^2+^ mobilisation and IP_1_ accumulation assays. By comparing to published and newly generated data for rat mGlu_5_ we show that while modulator pharmacology is relatively consistent across species, ligand dependent species differences in allosteric modulator affinity, cooperativity and probe dependence are evident. Additionally, we report PAM-dependent effects on orthosteric agonist kinetic profiles at human mGlu_5._ Together, these data highlight the importance of systematic evaluation of mGlu_5_ allosteric ligand activity at human mGlu_5_ to improve drug design and overcome potential barriers to translatability to clinical settings.

## 1. Introduction

Metabotropic glutamate receptor subtype 5 (mGlu_5_) is a Class C G protein-coupled receptor (GPCR) ubiquitously expressed throughout the mammalian CNS, playing an important role in both normal physiology and the pathophysiology of neurodegenerative and neuropsychiatric disorders (Gregory and Goudet, 2021). As such, mGlu_5_ has emerged as a promising drug target for multiple CNS disorders (Trinh *et al*., 2018; Budgett *et al*., 2022). However, targeting mGlu_5_ orthosteric ligand binding sites has been hampered due to extensive sequence conservation of orthosteric sites across mGlu subtypes, spurring the development of more selective agents, known as allosteric modulators (Sengmany and Gregory, 2016; Hellyer *et al*., 2017; Leach and Gregory, 2017). Allosteric modulators target less conserved sites topographically distinct from the orthosteric binding site and afford greater subtype selectivity. Allosteric ligands alter mGlu_5_ function through changes in orthosteric agonist affinity and/or efficacy, a property referred to as cooperativity (Christopoulos and Kenakin, 2002; Changeux and Christopoulos, 2016). In addition to improved selectivity, allosteric modulators offer additional advantages over orthosteric ligands (Melancon *et al*., 2012). First, pure allosteric modulators are quiescent in the absence of the endogenous agonist and will only modulate the receptor in the presence of an endogenous agonist, retaining spatial and temporal aspects of endogenous receptor signalling. Such ‘fine-tuning’ of physiological responses is an especially important consideration because with allosteric modulators receptor physiology is maintained, rather than silenced or completely switched on, potentially resulting in better therapeutic outcomes (Christopoulos and Kenakin, 2002; Leach *et al*., 2007; Melancon *et al*., 2012). Second, cooperativity between allosteric and orthosteric ligands is both reciprocal and saturable; allosteric modulators have a ceiling level to their effect, and may be safer than orthosteric ligands if administered in overdose (Lindsley *et al*., 2016).

Allosteric modulators which enhance or inhibit orthosteric ligand affinity and/or efficacy are positive allosteric modulators (PAMs) or negative allosteric modulators (NAMs), respectively (Changeux and Christopoulos, 2016). PAMs that activate the receptor in the absence of endogenous ligand are categorised as PAM-agonists or ago-PAMs. Additionally, neutral allosteric ligands (NALs) bind to allosteric sites but display no cooperativity with orthosteric ligands (Rodriguez *et al*., 2005; Melancon *et al*., 2012; Gregory *et al*., 2015). mGlu_5_ allosteric modulator discovery programs have discovered a broad spectrum of chemotypes including PAMs, NAMs and NALs (Lindsley *et al*., 2016; Gregory and Goudet, 2021). Multiple mGlu_5_ NAMs and NALs have advanced to clinical trials for anxiety, depression, Fragile X syndrome, Parkinson’s disease Levodopa-induced dyskinesias (PD-LID), gastroesophogeal reflux disorder (GERD) and migraine, although poor efficacy and safety have resulted in the termination of trials for many indications, and the outcomes of some are yet to be articulated. Additionally, mGlu_5_ PAMs have potential utility for treating schizophrenia and enhancing cognition (Emmitte, 2011; Gregory and Conn, 2015; Sengmany and Gregory, 2016; Gregory and Goudet, 2021). Despite extensive drug development programs, no mGlu_5_ targeted drugs have been marketed due to lack of efficacy, side effects or safety concerns in both preclinical and clinical settings (Porter *et al*., 2005; Jacob *et al*., 2009; Dekundy *et al*., 2011; Bridges *et al*., 2013; Hughes *et al*., 2013; Rook *et al*., 2013; Abou Farha *et al*., 2014; Swedberg and Raboisson, 2014; Swedberg *et al*., 2014). While it is clear mGlu_5_ allosteric modulation is a promising approach for treating neurological disorders, it is also evident there are limitations in current drug discovery paradigms resulting in optimisation of lead compounds that ultimately fail in preclinical and/or clinical trials.

Functional studies of mGlu_5_ allosteric modulators are predominantly performed using recombinantly expressed rat mGlu_5_ and pharmacological characterisation in native cell systems *in vitro* have been limited to non-human brain slice or neuronal cultures such as mouse cortical, striatal and hippocampal neurons and rat astrocytes (Gasparini *et al*., 1999; O’Brien *et al*., 2004; Noetzel *et al*., 2013; Rook *et al*., 2015; Balu *et al*., 2016; Sengmany *et al*., 2017, 2019; Hellyer *et al*., 2019). While some comparisons have been made between modulator pharmacology at human and rodent mGlu_5_, the use of different cell lines, experimental paradigms and analysis methods make comparisons across studies difficult (Lindsley *et al*., 2004; O’Brien *et al*., 2004; Kinney *et al*., 2005; Porter *et al*., 2005; Gichinga *et al*., 2011; Felts *et al*., 2013; Rook *et al*., 2013, 2015; Conde-Ceide *et al*., 2015). Modulator pharmacology is also largely tested using a single functional assay (Lindsley *et al*., 2016). However mGlu_5_ is pleiotropically coupled and signals through multiple intracellular pathways, with both PAMs and NAMs displaying biased pharmacology at rodent mGlu_5_, a phenomenon in which ligands display divergent pharmacology across distinct signalling pathways (Gregory *et al*., 2012; Kenakin and Christopoulos, 2013; Sengmany *et al*., 2017, 2019). In addition, differential pharmacology is evident between modulators that bind to multiple, distinct mGlu_5_ allosteric binding sites (O’Brien *et al*., 2004; Chen *et al*., 2008; Hammond *et al*., 2010). How such complexities apply to human mGlu_5_ is unknown. The lack of translatability from preclinical to clinical settings calls into question the ability to predict mGlu_5_ drug behaviour in humans based on non-human mGlu_5_ pharmacological data. As such, there remains a need to better quantify and assess the impact of chemically and pharmacologically diverse allosteric modulators on human mGlu_5_ activity.

In the current study, we rigorously characterised the pharmacology of eleven mGlu_5_ allosteric modulators across radioligand binding and distinct functional measures (iCa^2+^ mobilisation and inositol phosphate (IP_1_) accumulation) using HEK293A cells stably expressing human mGlu_5_. Allosteric ligands were representative of diverse chemical scaffolds and pharmacological profiles, including second-site PAMs, ago-PAMs, partial and full NAMs and NALs. Here, we report on species differences in the molecular pharmacological properties among the mGlu_5_ allosteric ligands tested. Differences in allosteric modulator affinity and cooperativity were evident depending on the agonist, species and method of analysis. These data highlight the importance of robust evaluation of mGlu_5_ allosteric ligand activity at human mGlu_5_ to better appreciate the full scope of ligand pharmacology and to shed light on potential barriers to translatability between preclinical and clinical settings.

## 2. Materials and Methods

### 2.1 Materials

Dulbecco’s modified Eagle’s medium (DMEM), and lipofectamine^TM^ 2000 were purchased from Invitrogen (Carlsbad, CA). Fetal bovine serum (FBS) was purchased from Thermo Fisher Scientific (Melbourne, Australia). Fluo8-AM and (S)-3,5-dihydroxyphenylglycine (DHPG) were purchased from Abcam (Cambridge, UK). Ionomycin and (4-fluorophenyl)(2-(phenoxymethyl)-6,7-dihydrooxazolo[5,4-c]pyridin-5(4H)-yl)methanone (VU0409551) were purchased from Sapphire Bioscience (Redfern, Australia). Select mGlu_5_ modulators 5-methyl-2-(phenylethynyl)pyridine (5MPEP), 3-cyano-*N*-(1,3-diphenyl-1*H*-pyrazol-5-yl) benzamide (CDPPB), 1-(4-(2,4-difluorophenyl)piperazin-1-yl)-2-((4-fluorobenzyl)oxy)ethanone (DPFE), N-(3-chloro-2-fluorophenyl)-3-cyano-5-fluorobenzamide (VU0366248), 3-fluoro-N-(4-methyl-2-thiazolyl)-5-(5-pyrimidinyloxy)benzamide (VU0409106) and (R)-5-((3-fluorophenyl) ethynyl)-N-(3-hydroxy-3-methylbutan-2-yl)picolinamide (VU0424465) were a gift from Vanderbilt University and synthesised as described previously (Kinney *et al*., 2005; Rodriguez *et al*., 2005; Felts *et al*., 2010, 2013; Gregory *et al*., 2013; Rook *et al*., 2013). N-(3-chlorophenyl)-3-[(2e)-1-methyl-4-oxoimidazo-lidin-2-ylidene]urea (fenobam) and 2-methyl-6-(phenylethynyl)pyridine (MPEP) were purchased from HelloBio (Bristol, UK). *N*-[4-chloro-2-[(1,3-dioxo-1,3-dihydro-2*H*-isoindol-2-yl)methyl]phenyl]-2-hydroxy-benzamide (CPPHA) was purchased from Focus Bioscience (Murrarie, Australia). 4-butoxy-*N*-(2,4-difluorophenyl)benzamide (VU0357121) was purchased from Tocris (Bristol, UK). [^3^H]methoxy-PEPy was custom synthesised by Pharmaron (Pharmaron, Inc. Beijing, China). Unless otherwise stated, all other reagents were purchased from Merck/Sigma-Aldrich (St. Louis, MO) and were of analytical grade.

### 2.2 Cell Culture

HEK293A cells were purchased from Invitrogen, were not used past passage 30 and were regularly screened for mycoplasma contamination. HEK293A cells stably expressing human mGlu_5_ (hereafter referred to as HEK293A-hmGlu_5_) were generated by transfecting HEK293A cells with human mGlu_5_ (alpha splice variant) DNA in pCDNA5 using lipofectamine^TM^ 2000 at a 1:6 ratio of DNA:lipofectamine. Human mGlu_5_ cells were selected by maintaining cells in humidified incubators at 37 °C and 5% CO_2_ in DMEM supplemented with 5% FBS, 16 mM HEPES and 500µg/mL geneticin for 5 passages. Polyclonal cell lines were then maintained in DMEM supplemented with 5% FBS, 16 mM HEPES. HEK293A cells stably expressing wild-type rat mGlu_5_ at low levels (hereafter referred to as HEK293A-rmGlu_5_) were generated previously and maintained under the same conditions (Noetzel *et al*., 2012). One day prior to experimentation, cells were plated onto poly-D-lysine coated, clear walled (iCa^2+^ mobilisation and IP_1_ accumulation) or white walled (radioligand binding) clear-bottom 96 well plates in glutamine-free DMEM supplemented with 5% dialysed FBS and 16 mM HEPES at a density of 40,000 cells/well.

### 2.3 Whole cell radioligand binding

For saturation binding studies, cells were incubated with increasing concentration of [^3^H]methoxy-PEPy on a shaker at RT for 1 hr in binding buffer (MgCl_2_.6H_2_O 0.49mM, MgSO_4_.7H_2_O 0.41mM, KCl 5.33 mM, KH_2_PO_4_ 0.44 mM, NaCl 137.93 mM, Na_2_HPO_4._7H_2_0 (dibasic) 0.34 mM, D-glucose 5.56 mM supplemented with 20 mM HEPES and 1.2 mM CaCl_2_, pH 7.4). For inhibition binding studies, cells were incubated with ∼3nM [^3^H]methoxy-PEPy in the presence of increasing concentrations of allosteric modulators on a shaker at RT for 1 hr in binding buffer. The DMSO concentration (0.3%) was kept constant throughout. Non-specific binding was determined in the presence of 10 μM MPEP. Assays were terminated by washing three times with ice-cold 0.9% NaCl and plates allowed to dry overnight. The next day UltimaGold (100 μl/well) was added directly to plates, which were sealed and incubated for >2h. Bound radioactivity was measured using a MicroBeta2 plate counter (PerkinElmer, Waltham, USA).

### 2.4 Intracellular calcium (iCa^2+^) mobilisation assay

All iCa^2+^ mobilisation assays were carried out in calcium assay buffer (MgCl_2_.6H_2_O 0.49mM, MgSO_4_.7H_2_O 0.41mM, KCl 5.33 mM, KH_2_PO_4_ 0.44 mM, NaCl 137.93 mM, Na_2_HPO_4._7H_2_0 (dibasic) 0.34 mM, D-glucose 5.56 mM supplemented with 20 mM HEPES, 1.2 mM CaCl_2_ and 4mM probenecid, pH 7.4). iCa^2+^ mobilisation was measured as a change in fluorescence of the cell permeable Ca^2+^ indicator dye Fluo-8AM at room temperature using Flexstation I and/or III (Molecular Devices) as described previously (Gregory *et al*., 2012). Baseline fluorescence of each individual well was determined for 20 sec prior to addition of agonists/modulators for all paradigms. For agonism, a single add paradigm was used, where agonists and allosteric modulators were added and fluorescence measured for a further 1 min. For modulation studies, a double add paradigm was used; allosteric ligands or vehicle were added 1 min prior to orthosteric agonist, with the exception of VU0424465 which was added simultaneously with orthosteric agonist to avoid the confounding effects of desensitisation (Hellyer *et al*., 2019). A 5-point smoothing function was applied to the raw fluorescence traces. Peak fluorescence was defined as the change from corresponding baseline and values were normalised to the maximal response to ionomycin (for agonism), glutamate or DHPG (for modulation). For area under curve (AUC) analysis, raw kinetic fluorescence traces were plotted, the mean of first 20 sec was defined as basal and the total peak area in response to different concentrations of either allosteric modulators, glutamate or DHPG (in absence or presence of modulators) over 60 sec post-addition were plotted using GraphPad Prism 10 (GraphPad, San Diego, CA).

### 2.5 Inositol monophosphate (IP_1_) accumulation assay

HEK293A-hmGlu_5_ cells were washed once and incubated for 1 hour at 37 °C with stimulation buffer (MgCl_2_.6H_2_O 0.49mM, MgSO_4_.7H_2_O 0.41mM, KCl 5.33 mM, KH_2_PO_4_ 0.44 mM, NaCl 137.93 mM, Na_2_HPO_4._7H_2_0 (dibasic) 0.34 mM, D-glucose 5.56 mM supplemented with 20mM HEPES, 30mM LiCl_2_, 1.2mM CaCl_2_, pH 7.4). Glutamic pyruvic transaminase (GPT; 10 U/ml) and sodium pyruvate (6 mM) were added to reduce ambient extracellular glutamate, which may confound observations of allosteric agonism (Sengmany et al., 2017). Compounds were diluted in stimulation buffer and incubated with cells for 1 hr at 37 °C prior to cell lysis with lysis buffer (50mM HEPES, 15mM potassium fluoride, 1.5% Triton X-100, 3% FBS and 0.2% BSA, pH 7) for 30 min at room temperature. IP_1_ levels were determined using the HTRF® IP-one assay kit as per manufacturer’s instructions and fluorescence measured at 615 and 665 nm using the Envision plate reader (PerkinElmer) after excitation at 340 nm. Data is expressed as fold of basal IP_1_ accumulation.

### 2.6 Data analysis

All nonlinear regression analyses were performed using Prism 9 (GraphPad Software Inc, San Diego, CA). In most cases, curve fitting settings were kept default, with least squares regression, medium convergence with a maximum iteration number of 1000, no weighting and no special handling of outliers (Motulsky and Brown, 2006). Saturation binding curves were analysed by simultaneously fitting total and non-specific binding curves to equation 1:

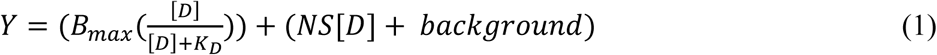

Where B_max_ is the maximum specific binding of [^3^H]methoxy-PEPy, the molar concentration of [^3^H]methoxy-PEPy is [D] and K_D_ is the radioligand equilibrium dissociation constant, NS is the slope of linear nonspecific binding and background is the background signal when no radioligand is present.

Inhibition of [^3^H]methoxy-PEPy binding data were fitted to either an allosteric modulator titration model (equation 2) or either a one-site or two-site inhibition binding model as previously described (Gregory et al., 2012; Lazareno and Birdsall, 1995). An extra sum-of-squares F test was used to determine the preferred model for each data set.

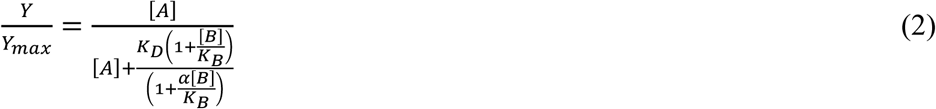

Where Y/Y_max_ is the fractional specific binding, [A] is the radioligand concentration, [B] is the concentration of the allosteric modulator, K_D_ is the radioligand equilibrium dissociation constant, K_B_ is the allosteric modulator equilibrium dissociation constant and α is the binding cooperativity factor. An α value of α > 1 denotes positive cooperativity, values of 0 > α < 1 denote negative cooperativity, and α=1 denotes neutral cooperativity. K_D_ was constrained to values determined from [^3^H]methoxy-PEPy saturation binding experiments (see Table 1).

**Table 1.**
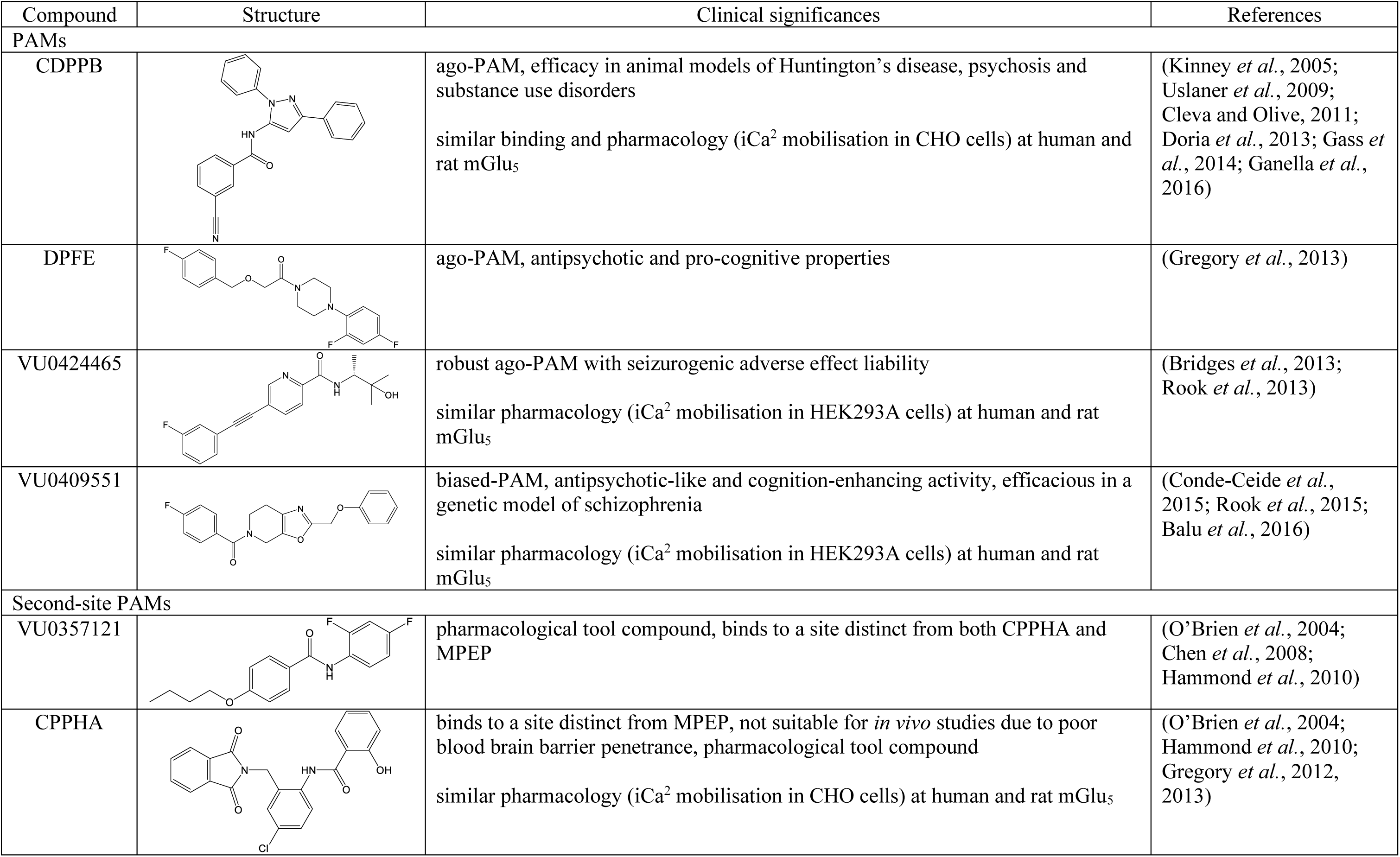
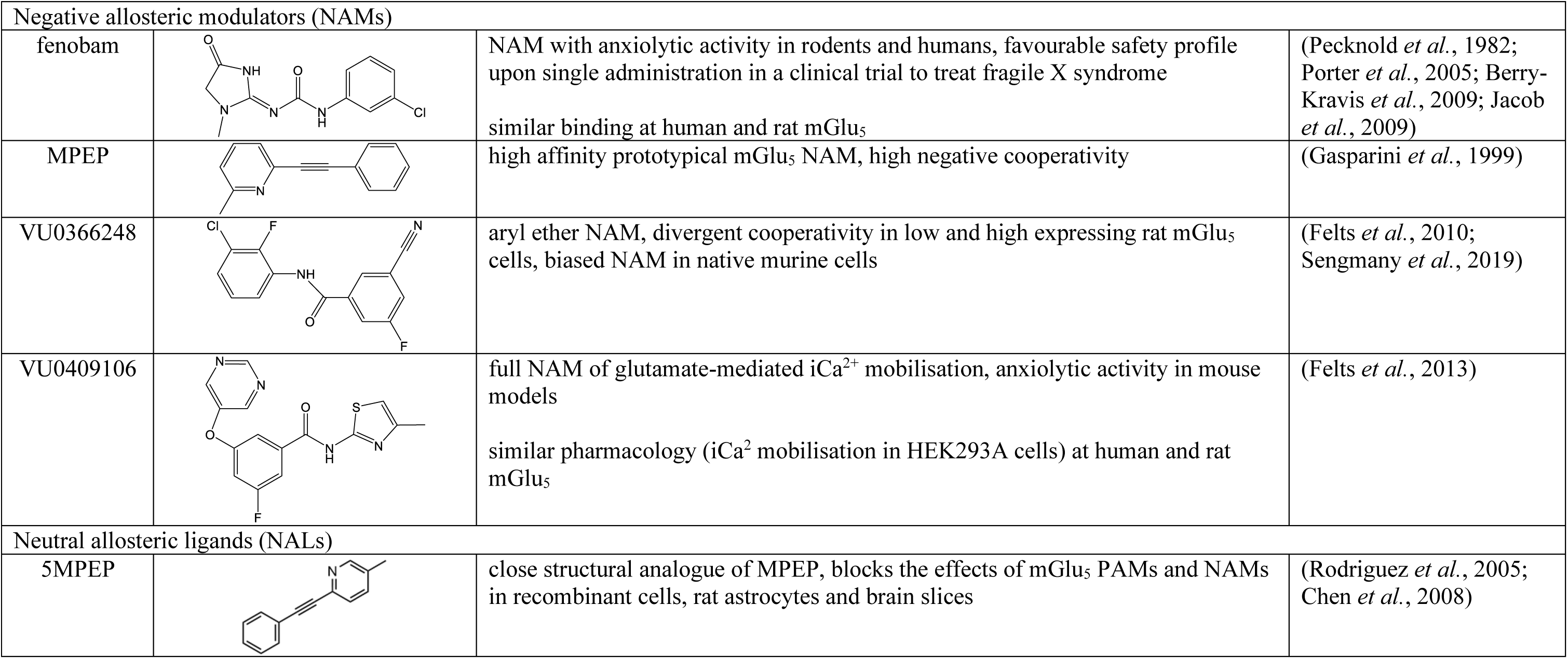
Structures, clinical significance and previously reported species-dependent pharmacology of allosteric modulators previously classified as either PAMs, second-site PAMs, NALs or NAMs investigated in the current study.

| Compound | Structure | Clinical significances | References |
| --- | --- | --- | --- |
| **PAMs** |  |  |  |
| CDPPB |  | ago-PAM, efficacy in animal models of Huntington's disease, psychosis and substance use disorders<br>similar binding and pharmacology (iCa<sup>2</sup> mobilisation in CHO cells) at human and rat mGlu<sub>5</sub> | (Kinney *et al.*, 2005; Uslaner *et al.*, 2009; Cleva and Olive, 2011; Doria *et al.*, 2013; Gass *et al.*, 2014; Ganella *et al.*, 2016) |
| DPFE |  | ago-PAM, antipsychotic and pro-cognitive properties | (Gregory *et al.*, 2013) |
| VU0424465 |  | robust ago-PAM with seizurogenic adverse effect liability<br>similar pharmacology (iCa<sup>2</sup> mobilisation in HEK293A cells) at human and rat mGlu<sub>5</sub> | (Bridges *et al.*, 2013; Rook *et al.*, 2013) |
| VU0409551 |  | biased-PAM, antipsychotic-like and cognition-enhancing activity, efficacious in a genetic model of schizophrenia<br>similar pharmacology (iCa<sup>2</sup> mobilisation in HEK293A cells) at human and rat mGlu<sub>5</sub> | (Conde-Ceide *et al.*, 2015; Rook *et al.*, 2015; Balu *et al.*, 2016) |
| **Second-site PAMs** |  |  |  |
| VU0357121 |  | pharmacological tool compound, binds to a site distinct from both CPPHA and MPEP | (O'Brien *et al.*, 2004; Chen *et al.*, 2008; Hammond *et al.*, 2010) |
| CPPHA |  | binds to a site distinct from MPEP, not suitable for *in vivo* studies due to poor blood brain barrier penetrance, pharmacological tool compound<br>similar pharmacology (iCa<sup>2</sup> mobilisation in CHO cells) at human and rat mGlu<sub>5</sub> | (O'Brien *et al.*, 2004; Hammond *et al.*, 2010; Gregory *et al.*, 2012, 2013) |

| Negative allosteric modulators (NAMs) |  |  |  |
| --- | --- | --- | --- |
| fenobam |  | NAM with anxiolytic activity in rodents and humans, favourable safety profile upon single administration in a clinical trial to treat fragile X syndrome<br>similar binding at human and rat mGlu<sub>5</sub> | (Pecknold *et al.*, 1982; Porter *et al.*, 2005; Berry-Kravis *et al.*, 2009; Jacob *et al.*, 2009) |
| MPEP |  | high affinity prototypical mGlu<sub>5</sub> NAM, high negative cooperativity | (Gasparini *et al.*, 1999) |
| VU0366248 |  | aryl ether NAM, divergent cooperativity in low and high expressing rat mGlu<sub>5</sub> cells, biased NAM in native murine cells | (Felts *et al.*, 2010; Sengmany *et al.*, 2019) |
| VU0409106 |  | full NAM of glutamate-mediated iCa<sup>2+</sup> mobilisation, anxiolytic activity in mouse models<br>similar pharmacology (iCa<sup>2+</sup> mobilisation in HEK293A cells) at human and rat mGlu<sub>5</sub> | (Felts *et al.*, 2013) |
| Neutral allosteric ligands (NALs) |  |  |  |
| 5MPEP |  | close structural analogue of MPEP, blocks the effects of mGlu<sub>5</sub> PAMs and NAMs in recombinant cells, rat astrocytes and brain slices | (Rodriguez *et al.*, 2005; Chen *et al.*, 2008) |

Agonist-concentration response curves were fitted to a variable four-parameter logistic equation 3:

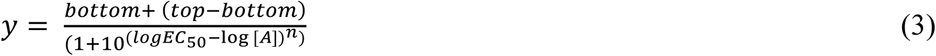

Where bottom and top are lower and upper plateau levels of the concentration response curve respectively, *n* is the Hill coefficient, [A] is the log molar concentration of agonist, and EC_50_ is the agonist concentration required to produce a half maximal response between top and bottom values (potency).

Partial agonist concentration response curves were generated by fitting an operational model of partial agonism, where full agonist concentration-response relationships were fitted to equation 3, and partial agonists were simultaneously fitted to equation 4:

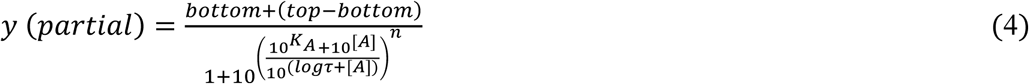

where bottom, top, [A], EC_50_ and *n* are as described above. K_A_ is the affinity of the partial agonist, and logτ is the transducer constant describing the efficacy of the partial agonist.

Orthosteric agonist concentration-response curves in the absence and presence of increasing concentrations of allosteric modulators were fitted to a complete operational model of allosterism (equation 5) to estimate affinity (K_B_), cooperativity (αβ) and efficacy (τ_B_) (Leach *et al*., 2007):

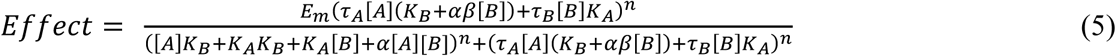

where [A] and [B] are the log molar concentration of orthosteric agonist and allosteric modulator respectively. K_A_ and K_B_ are the equilibrium dissociation constants of the orthosteric agonist and allosteric modulator respectively. E_m_ is the maximal system response, τ_A_ and τ_B_ are the efficacy of orthosteric and allosteric ligands, respectively, α and β denote allosteric effects on orthosteric ligand-binding affinity and efficacy, respectively. Where possible, τ_B_ was constrained to estimates derived from partial agonist equations for PAM-agonists, and was constrained to −100 (i.e. no allosteric agonism) for pure PAMs and NAMs. Affinity cooperativity (α) was assumed to be neutral as validated previously with rat mGlu_5_ (Gregory et al., 2012) and thus constrained to a value of 1, allowing estimates of β as a measure of cooperativity. For full NAMs that completely inhibit the orthosteric ligand responses, such that β approaches to zero, logβ was constrained to a value of −100 to allow curve fitting and estimation of affinity. pK_A_ values were constrained to affinity estimates previously determined from inhibition binding assays for glutamate (6.16 and 5.45 for rat and human mGlu_5_, respectively) and DHPG (5.41 for rat mGlu_5_) (Mutel *et al*., 2002; Ohashi *et al*., 2002). K_A_ estimates for DHPG at human mGlu_5_ were determined by using an irreversible allosteric ligand (RVDU-3-185) to covalently bind and functionally deplete receptors, followed by iCa^2+^ mobilisation assays with increasing concentrations of DHPG and fitting to a model of receptor depletion (equation 6) to derive affinity and efficacy estimates (Gregory *et al*., 2016).

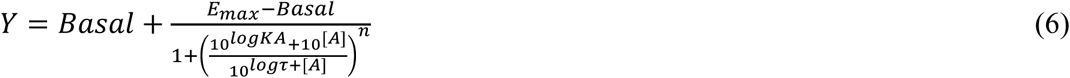

Where [A], K_A_, E_max_ and n have described earlier, τ is the transducer constant, an index of agonist efficacy. The pK_A_ estimate derived for DHPG at human mGlu_5_ using this method was 5.78 ± 0.18.

Time course data for orthosteric agonist-induced iCa^2+^ mobilisation were fitted to the empirical time course equation “rise-and-fall to baseline” (equation 7) (Hoare, Tewson, Quinn, Hughes, *et al*., 2020).

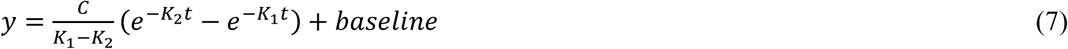

where C is the initial rate of signalling, defined as a constant (y-units.t^−1^) and K_1_ and K_2_ are observed rate constants of the rise phase and fall phase, respectively, in units of t^−1^. K_1_ is constrained to be the larger of the two rate constant values (i.e. > K_2_).

All parameters were estimated and represented as logarithmic mean ± SEM. Statistical analyses of saturation binding was performed using a student’s t-test to compare K_D_, B_max_ and cooperativity estimates between rat and human mGlu_5_. Statistical analyses of potency and affinity estimates from inhibition binding and functional assays were performed using one-way analysis of variance (ANOVA) with Sidak’s multiple comparison to compare affinity and cooperativity estimates between assays. Statistical analyses of kinetic parameter estimates were performed using extra sum-of-squares F-test to compare kinetic fits in the presence and absence of different PAMs. Statistical significance was defined as p < 0.05 of a group of 3-7 independent experiments performed in duplicate.

## 3. Results

Eleven structurally diverse allosteric modulators previously categorised as PAMs, ago-PAMs, second-site PAMs, NAMs and NALs based on glutamate stimulation of rat-mGlu_5_ mediated iCa^2+^ mobilisation assays were selected for the study (Table 1). To compare modulator binding and function between rat and human mGlu_5_, comparisons were primarily made to published data from our laboratory where possible (i.e. where binding and functional experiments had been conducted for rat mGlu_5_ under the same conditions) as differences between cell lines, experimental paradigms and data analysis precluded comparisons from independent sources. For functional studies, multiple estimates for rat mGlu_5_ were considered when derived from assays performed in the same cell lines, under the same conditions and with the same analyses applied. Where no data were available, binding or functional studies were performed in HEK293A-rat mGlu_5_ cells as noted throughout.

### 3.1. Binding affinity and cooperativity for diverse mGlu_5_ allosteric ligands in HEK293A-hmGlu_5_ cells

The radioligand [^3^H]methoxy-PEPy is structurally similar to MPEP and binds selectively at the common allosteric site, also referred to as “MPEP-site”, within the mGlu_5_ 7TM bundle (Cosford *et al*., 2002). Whole-cell [^3^H]methoxy-PEPy saturation radioligand binding experiments were initially conducted to characterise radioligand binding properties and receptor expression in the newly generated HEK293A-hmGlu_5_ cell line prior to inhibition binding studies. [^3^H]methoxy-PEPy affinity was significantly higher at human compared to rat mGlu_5_ (pK_D_ of 8.34 ± 0.05 vs 7.78 ± 0.10, respectively; p < 0.05), but B_max_ was similar across cell lines (14.1 ± 0.8 fmol/10^4^ cells and 14.9 ± 4.5 fmol/10^4^ cells for hmGlu_5_ and rmGlu_5_, respectively).

Whole-cell [^3^H]methoxy-PEPy inhibition radioligand binding experiments were then conducted to determine binding affinities of mGlu_5_ allosteric modulators at hmGlu_5_. Of all modulators, only VU0409551, DPFE, 5MPEP concentration-response relationships were best fitted to a one-site competitive binding model (Figure 1). Additionally, MPEP displayed biphasic binding, with inhibition binding data best fitted to a two-site binding model (Figure 1, Table 2). For all other modulators, [^3^H]methoxy-PEPy, inhibition binding data were best fitted to an allosteric modulator titration model, which was used to derive affinity (pK_B_) and cooperativity (logα) estimates (Figure 1, Table 2). The rank order of PAM affinity for hmGlu_5_ was VU0424465 > CDPPB > VU0357121 > VU0409551 > CPPHA ≥ DPFE. The rank order of NAM affinity for hmGlu_5_ was MPEP (high affinity site) > fenobam > VU0409106 > VU0366248 ≥ MPEP (low affinity site) (Table 2). The cooperativity of VU0366248, CDPPB, VU0357121 and CPPHA with [^3^H]methoxy-PEPy were all similar, reflecting the ability of these ligands to only displace ∼50% of [^3^H]methoxy-PEPy binding and indicative of non-competitive binding when measured in whole cells. While VU0424465, VU0409106 and fenobam inhibition binding data were all best fitted to an allosteric model, these compounds displayed higher negative cooperativity, displacing 80-95% of [^3^H]methoxy-PEPy binding (Figure 1, Table 2). The overall rank order of cooperativity was fenobam > VU0409106 = VU0424465 > VU0366248 ≥ CPPHA > CDPPB ≥ VU0357121 (Table 2).

**Figure 1.**
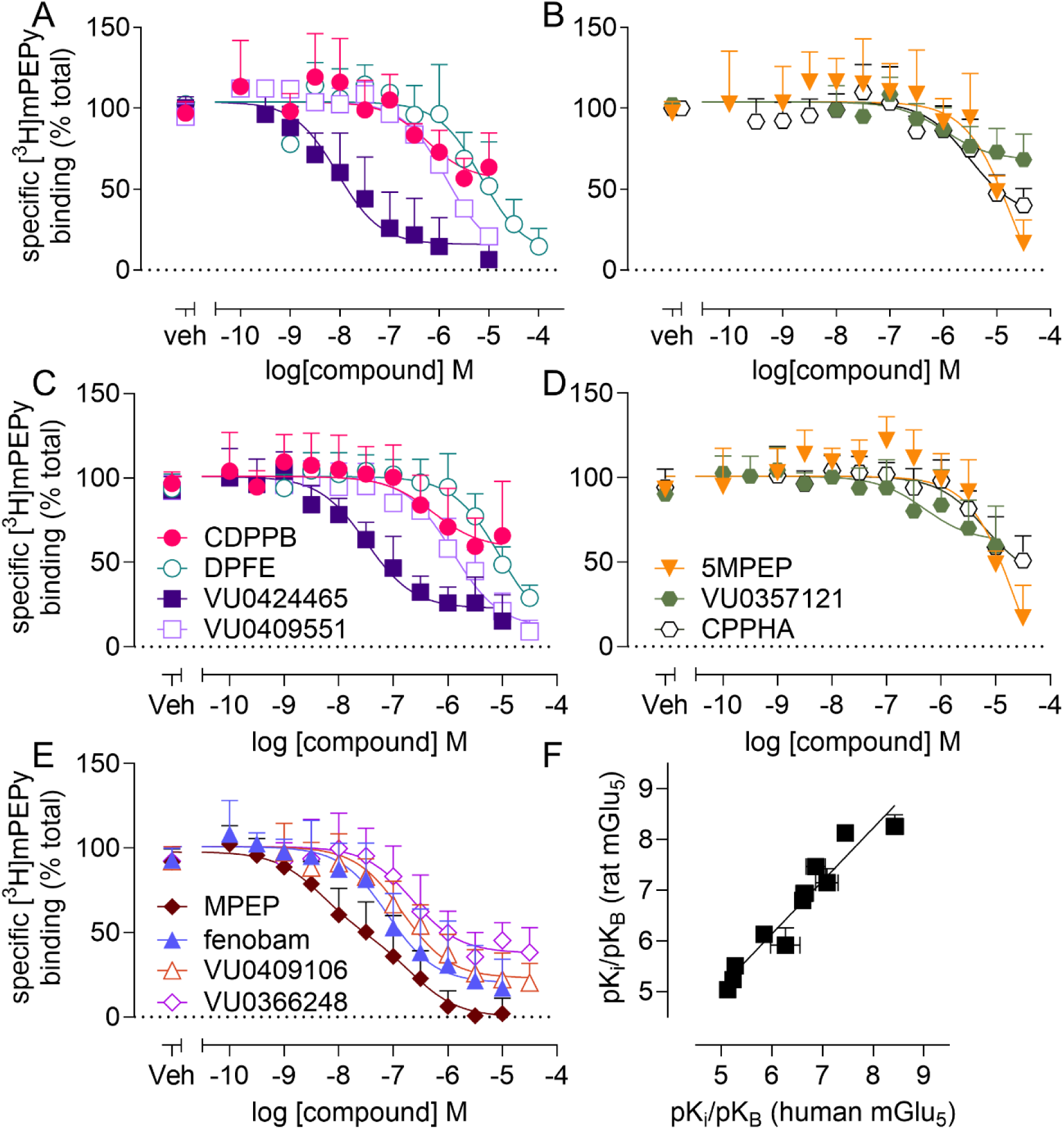
Inhibition of [^3^H]methoxy-PEPy binding to rat and human mGlu_5_ in whole cells by mGlu_5_ allosteric modulators. Inhibition of [^3^H]methoxy-PEPy binding was determined in intact, adherent HEK293A-rmGlu_5_ (**A, B**)or HEK293A-hmGlu_5_ (**C-E**) cells for allosteric modulators previously classified as PAMs (**A, C**), second-site PAMs and NAL (**B, D**) or NAMs (**E**). Data were fitted to either allosteric or competitive model to estimate affinity (pK_B_ and/or pK_i_) and cooperativity (logα) as determined by an F-test. (**F**) Correlation analysis reveals affinity values derived for allosteric modulator binding to rat mGlu_5_ highly correlate with those derived for binding to human mGlu_5_ (slope = 1.05 ± 0.10; r^2^ = 0.92). Data are mean + S.D. from at least three independent determinations performed in duplicate. The number of independent experiments performed in duplicate for each ligand are listed in Table 1.

**Table 2.** Summary of affinity (pK_B_ and/or pK_i_) and cooperativity (logα) estimates for mGlu_5_ allosteric modulators derived from [^3^H]methoxy-PEPy inhibition binding assays in HEK293A-hmGlu_5_ or HEK293A-rmGlu_5_ whole cells. Data represent mean ± S.E.M. from the indicated number of independent determinations performed in duplicate.

| | HEK293A-hmGlu<sub>5</sub> $pK_i/pK_B$<sup>a</sup> | | | | | HEK293A-rmGlu<sub>5</sub>-low $pK_i/pK_B$ | | | | |
| --- | --- | --- | --- | --- | --- | --- | --- | --- | --- | --- |
| | High<sup>b</sup> | Low<sup>c</sup> | Fraction High<sup>d</sup> | $\log\alpha$<sup>e</sup> | n | High | Low | Fraction High | $\log\alpha$ | n |
| VU0424465 | 7.45 ± 0.14* | N.D. | 1 | -0.86 ± 0.13 | 6 | 8.13 ± 0.13 | N.D. | 1 | -0.99 ± 0.14 | 4 |
| CDPPB | 6.61 ± 0.16 | N.D. | 1 | -0.32 ± 0.07 | 10 | 6.80 ± 0.14 | N.D. | 1 | -0.27 ± 0.04 | 5 |
| VU0409551 | 5.85 ± 0.11 | N.D. | 1 | N.A. | 7 | 6.13 ± 0.08 | N.D. | 1 | N.A. | 4 |
| DPFE | 5.13 ± 0.09 | N.D. | 1 | N.A. | 5 | 5.04 ± 0.12 | N.D. | 1 | N.A. | 3 |
| CPPHA | 5.28 ± 0.14 | N.D. | 1 | -0.50 ± 0.09 | 6 | 5.51 ± 0.07 | N.D. | 1 | -0.69 ± 0.13 | 7 |
| VU0357121 | 6.27 ± 0.29 | N.D. | 1 | -0.34 ± 0.10 | 5 | 5.92 ± 0.34 | N.D. | 1 | -0.39 ± 0.19 | 4 |
| 5MPEP | 5.24 ± 0.07 | N.D. | 1 | N.A. | 5 | 5.24 ± 0.07 | N.D. | 1 | N.A. | 5 |
| MPEP | 8.43 ± 0.17 | 6.48±0.26 | 0.50±0.12 | N.A. | 7 | 8.26 ± 0.23<sup>f</sup> | 6.25 ± 0.11<sup>f</sup> | 0.62 ± 0.11<sup>f</sup> | N.A. | 4 |
| fenobam | 7.09 ± 0.22 | N.D. | 1 | -1.06 ± 0.10 | 9 | 7.15 ± 0.28<sup>f</sup> | 4.66 ± 0.22<sup>f</sup> | 0.73 ± 0.08<sup>f</sup> | N.A. | 5 |
| VU0409106 | 6.87 ± 0.19* | N.D. | 1 | -0.88 ± 0.15 | 6 | 7.47 ± 0.07<sup>f</sup> | N.D. | 1 | -1.12 ± 0.12 | 4 |
| VU0366248 | 6.65 ± 0.17 | N.D. | 1 | -0.53 ± 0.07* | 6 | 6.94 ± 0.13<sup>f</sup> | N.D. | 1 | -0.77 ± 0.06 | 4 |

N.D. not determined due to allosteric model fit
N.A. not applicable due to one-site or two-site fit
<sup>a</sup> $pK_i/pK_B$ , negative logarithm of the equilibrium dissociation constant
<sup>b</sup> equilibrium dissociation constant for one-site binding fits or the high affinity site of two-site fits
<sup>c</sup> equilibrium dissociation constant for the low affinity site of two-site fits
<sup>d</sup> fraction of binding sites correlating to the high affinity site from two-site fits
<sup>e</sup> $\log\alpha$ , logarithm of the cooperativity factor for the interaction between the indicated allosteric modulator and [<sup>3</sup>H]methoxy-PEPy, derived from fitting inhibition binding data to an allosteric modulator titration model.
<sup>f</sup> Affinity and cooperativity estimates for MPEP, fenobam, VU0409106 and VU0366248 are from Sengmany *et al.*, 2019
\* significantly different ( $p < 0.05$ ) from estimates derived from HEK293A-rmGlu<sub>5</sub>, with one-way analysis of variance and Sidak's post hoc test

Whole cell [^3^H]methoxy-PEPy inhibition radioligand binding experiments were also conducted in HEK293A-rmGlu_5_ to determine binding affinities of mGlu_5_ allosteric modulators at rat mGlu_5_ where no equivalent literature values existed (for PAMs, second-site PAMs and NALs; Figure 1). Similar to hmGlu_5,_ VU0409551, DPFE, 5MPEP inhibition binding data at rat mGlu_5_ were best fitted to a one-site competitive binding model and MPEP displayed two-site binding (Table 2). In contrast to hmGlu_5_, fenobam displayed two-site binding at rmGlu_5_ (Sengmany *et al*., 2019). The rank order of PAM affinity for rmGlu_5_ was similar to hmGlu_5_, with the exception of VU0409551 displaying higher affinity than VU0357121. Similarly, VU0409106 had higher affinity than fenobam at rmGlu_5,_ to give a rank order of NAM affinity of MPEP (high affinity site) > VU0409106 > fenobam > VU0366248 > MPEP (low affinity site) > fenobam (low affinity site). When comparing across species, affinity values were mostly similar (Figure 1D), with the exception of VU0424465 and VU0409106, which had significantly higher affinity at rat compared to human mGlu_5,_ by 5-fold and 4-fold, respectively (Table 2). Cooperativity values were also mostly similar across species, with the exception of VU0366248, which had a 2-fold higher cooperativity estimate at rat mGlu_5_ compared to human mGlu_5._

### 3.2. mGlu_5_ allosteric modulators display differential allosteric agonism across functional readouts in HEK293A-hmGlu_5_ cells

Basic functional characterisation of the HEK293A-hmGlu_5_ cell line was initially performed using intracellular calcium (iCa^2+^) mobilisation assays. Two orthosteric agonists, the endogenous agonist glutamate and the non-membrane permeable surrogate agonist DHPG were initially tested (Figure 2A). Both orthosteric ligands robustly induced iCa^2+^ mobilisation in HEK293A-hmGlu_5_ cells. Relative to glutamate, DHPG was less potent and acted as a partial agonist, inducing ∼80% of maximal iCa^2+^ mobilisation. When tested in the absence of orthosteric agonist, VU0424465 displayed robust agonist activity in iCa^2+^ mobilisation assays, acting as a partial agonist relative to glutamate (Figure 2A). Glutamate (5.5-fold), DHPG (2.5-fold) and VU0424465 (3-fold) all had significantly lower potency estimates at human mGlu_5_ when compared to rat mGlu_5_ (Table 3). No other PAM or NAM had appreciable agonism or inverse agonist activity in the iCa^2+^ mobilisation assay (Figure 2).

**Figure 2.**
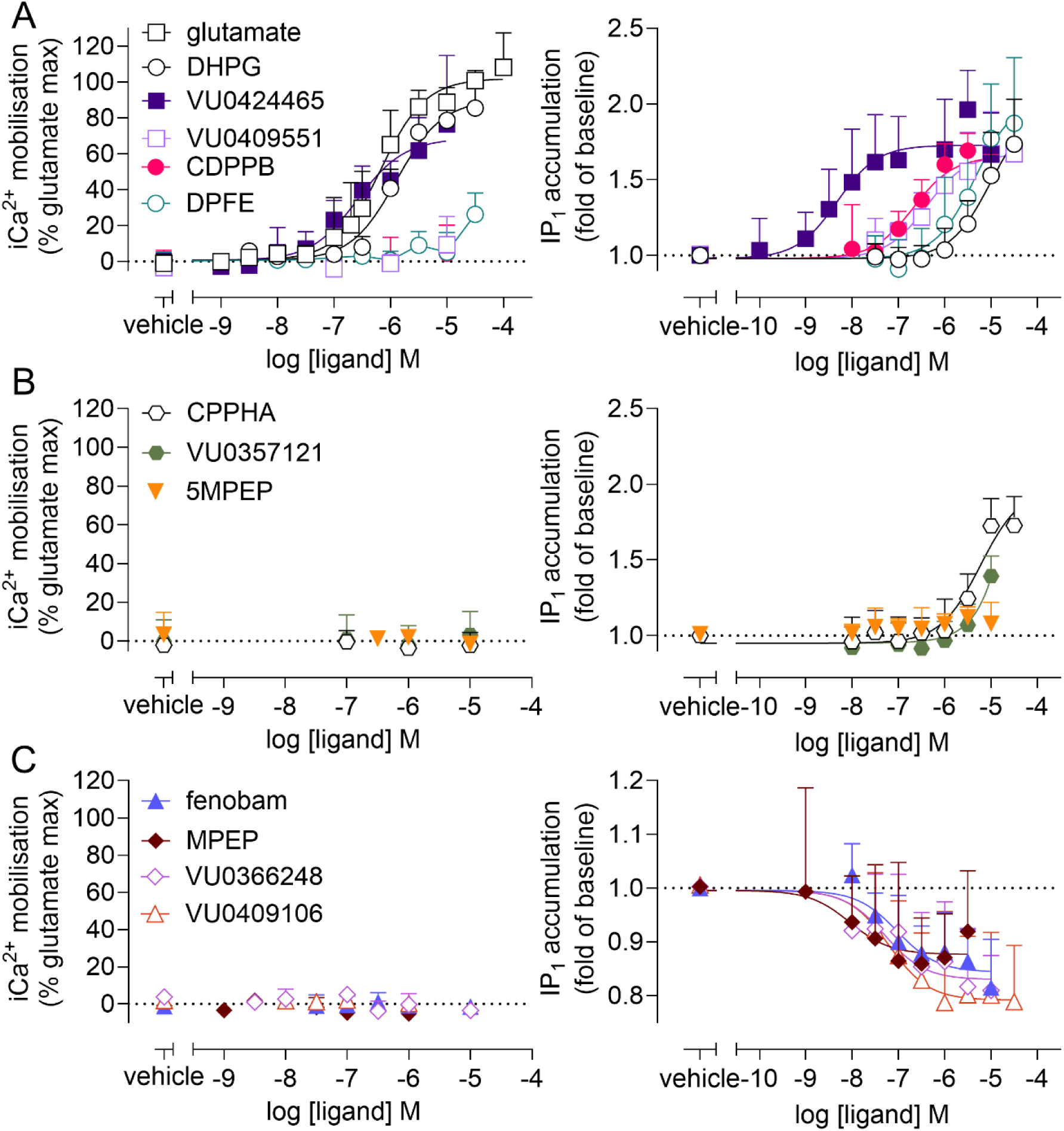
Agonist activity of mGlu_5_ orthosteric and allosteric ligands in iCa^2+^ mobilisation (left panels) and IP_1_ accumulation (right panels) assays in HEK293A-hmGlu_5_ cells. (**A**) orthosteric agonists and the ago-PAM VU0424465 induce iCa^2+^ mobilisation and IP_1_ accumulation. Other PAMs only induce IP_1_ accumulation. (**B**) Second-site PAMs and the NAL (5MPEP) have no activity in iCa^2+^ accumulation assays, but the second-site PAMs CPPHA and VU0357121 have agonist activity for IP_1_ accumulation. (**C**) NAMs do not have activity in iCa^2+^ mobilisation assays but induce inverse agonism in IP_1_ accumulation assays. Data are expressed as mean + S.D. from at least three independent determinations performed in duplicate. The number of independent experiments performed in duplicate for each ligand are denoted in Table 2. Error bars not shown lie within the dimensions of the symbols.

**Table 3.** Potency (pEC_50_) and maximum response estimates of orthosteric and allosteric agonism/inverse agonism of iCa^2+^ mobilisation and IP_1_ accumulation in HEK293A-hmGlu_5_ or HEK293A-rmGlu_5_ cells. Data represent mean ±SEM of the indicated number of independent experiments performed in duplicate.

|  | iCa<sup>2+</sup> mobilisation |  |  |  |  |  | IP<sub>1</sub> accumulation |  |  |  |  |  |
| --- | --- | --- | --- | --- | --- | --- | --- | --- | --- | --- | --- | --- |
|  | human mGlu<sub>5</sub> |  |  | rat mGlu<sub>5</sub> |  |  | human mGlu<sub>5</sub> |  |  | rat mGlu<sub>5</sub> |  |  |
|  | pEC<sub>50</sub><sup>a</sup> | E<sub>max</sub><sup>b</sup> | N | pEC<sub>50</sub> | E<sub>max</sub> | N | pEC<sub>50</sub>/pIC<sub>50</sub><sup>a</sup> | E<sub>max</sub>/I<sub>max</sub><sup>b</sup> | N | pEC<sub>50</sub>/pIC<sub>50</sub> | E<sub>max</sub>/I<sub>max</sub> | N |
| glutamate | 6.14 ± 0.08 | 103.4 ± 3.9 | 8 | 6.64 ± 0.06*<br>6.87 ± 0.06* | 106.4 ± 0.7<br>100.3 ± 0.2 | 7<sup>c</sup><br>26<sup>d</sup> | n.t. | n.t. | n.t. | n.t. | n.t. | n.t. |
| DHPG | 5.96 ± 0.03 | 80.3 ± 2.5 | 8 | 6.34 ± 0.05*<br>6.32 ± 0.08* | 102.6 ± 1.0<br>92.9 ± 2.6 | 8<sup>c</sup><br>10<sup>d</sup> | n.d. | 1.7 ± 0.11 | 7 | 5.68 ± 0.17 | 1.9 ± 0.20 | 6<sup>d</sup> |
| VU0424465 | 6.56 ± 0.12 | 69.5 ± 4.5 | 8 | 7.12 ± 0.14<sup>^</sup> | 74.8 ± 8.4 | 6<sup>d</sup> | 8.13 ± 0.14<sup>#</sup> | 1.7 ± 0.14 | 5 | 8.70 ± 0.13<sup>^</sup> | 1.9 ± 0.20 | 4<sup>d</sup> |
| CDPPB | n.a. | n.a. | 8 | 7.35 ± 0.21 | 18.9 ± 4.9 | 5<sup>d</sup> | 6.67 ± 0.11 | 1.7 ± 0.06 | 5 | 7.60 ± 0.18* | 2.3 ± 0.10* | 4<sup>d</sup> |
| VU0409551 | n.a. | n.a. | 5 | 6.23 ± 0.15 | 55.3 ± 5.8 | 4<sup>d</sup> | 6.18 ± 0.11 | 1.6 ± 0.14 | 5 | 6.16 ± 0.13 | 1.7 ± 0.20 | 3<sup>d</sup> |
| DPFE | n.a. | n.a. | 8 | 5.72 ± 0.17 | 49.7 ± 8.7 | 9<sup>d</sup> | n.d. | 1.8 ± 0.22 | 4 | 5.95 ± 0.12 | 2.7 ± 0.20* | 4<sup>d</sup> |
| CPPHA | n.a. | n.a. | 5 | n.d. | n.d. | 5<sup>c</sup> | n.d. | 1.9 ± 0.09 | 4 | n.r. | n.r. | 5<sup>c</sup> |
| VU0357121 | n.a. | n.a. | 5 | n.a. | n.a. | 5<sup>c</sup> | n.d. | 1.4 ± 0.05 | 6 | n.d. | 2.0 ± 0.16 | 5<sup>c</sup> |
| MPEP | n.a. | n.a. | 3 | n.a. | n.a. | 5<sup>e</sup> | 8.05 ± 0.30 | 0.87 ± 0.04 | 5 | 7.86 ± 0.27 | 0.74 ± 0.03* | 8<sup>e</sup> |
| fenobam | n.a. | n.a. | 5 | n.a. | n.a. | 5<sup>e</sup> | 7.26 ± 0.11 | 0.85 ± 0.02 | 5 | 6.83 ± 0.26 | 0.76 ± 0.03 | 9<sup>e</sup> |
| VU0366248 | n.a. | n.a. | 3 | n.a. | n.a. | 5<sup>e</sup> | 6.94 ± 0.52 | 0.84 ± 0.03 | 4 | 6.48 ± 0.16 | 0.81 ± 0.05 | 7<sup>e</sup> |
| VU0409106 | n.a. | n.a. | 5 | n.a. | n.a. | 5<sup>e</sup> | 7.46 ± 0.28 | 0.82 ± 0.04 | 5 | 6.89 ± 0.26 | 0.74 ± 0.05 | 7<sup>e</sup> |

NB: 5MPEP (NAL) had no agonist activity across any functional measures for either species (n=5-8)
n.t. glutamate was not tested due to the presence of GPT in assay buffer
n.r. not recorded as no data is available for rat mGlu<sub>5</sub>
n.a. not applicable as no agonism was evident.
n.d. not determined due to poor curve fit
<sup>a</sup> Negative logarithm of the concentration of agonist required to generate the half-maximal or inhibitory response derived from a three-parameter logistic equation.
<sup>b</sup> Maximal response to indicated modulators expressed as percentage of maximum glutamate response (iCa<sup>2+</sup> mobilisation) or fold-change from basal (IP<sub>1</sub> accumulation).
<sup>c</sup> data with rat mGlu<sub>5</sub> generated for this study
<sup>d</sup> data with rat-mGlu<sub>5</sub> retrieved from Sengmany *et al.*, 2017
<sup>e</sup> data with rat-mGlu<sub>5</sub> retrieved from Sengmany *et al.*, 2019

Previous studies revealed mGlu_5_ allosteric modulators display biased agonism/inverse agonism i.e. differential agonist activity across functional endpoints and kinetic conditions, with enhanced activity in IP_1_ accumulation assays (Sengmany *et al*., 2017, 2019). As such, modulators were next tested for agonist activity at human mGlu_5_ using IP_1_ accumulation assays (Figure 2). As IP_1_ accumulation is assessed over a longer period than iCa^2+^ mobilisation (1 hr vs 2 min), glutamic pyruvic transaminase (GPT) was added to IP_1_ assay buffer to reduce the effect of any ambient extracellular glutamate produced by cells, which may confound observations of allosteric agonism. Inclusion of GPT precluded the use of glutamate in these assays, and only DHPG was tested as an orthosteric agonist. DHPG, all PAMs and second-site PAMs had intrinsic agonist activity in IP_1_ accumulation assays using human mGlu_5_. DHPG was less potent for IP_1_ accumulation relative to iCa^2+^ mobilisation, with concentration response curves not reaching a maximal plateau and precluding accurate potency estimation (Figure 2A). VU0424465, VU0409551, CDPPB, DPFE and CPPHA all achieved a similar maximal response to DHPG, whereas VU0357121 achieved ∼60% maximal response compared to other agonists (Figure 2A & B). 5MPEP had no agonist activity in IP_1_ accumulation assays (Figure 2B). All NAMs displayed inverse agonist activity, reducing levels of IP_1_ to 0.8-0.9-fold of basal (Figure 2C), mirroring our previous observations at rat mGlu_5_ (Sengmany *et al.,* 2019).

Similar to DHPG, concentration response curves for DPFE, CPPHA and VU0357121 did not reach plateau over the tested concentration ranges; potency estimates could therefore not be derived and E_max_ estimates are given as the IP_1_ response at the highest concentration tested. However, potency and E_max_ estimates could be derived for VU0424465, CDPPB, VU0409551, and all NAMs. Relative to iCa^2+^ mobilisation, VU0424465 had a significantly higher potency (30-fold) for IP_1_ accumulation at human mGlu_5_ (Table 3). Compared to rat mGlu_5_, VU0409551 and all NAMs had similar potency estimates, whereas VU0424465 and CDPPB had significantly lower potency estimates, by 4-fold and 8.5-fold, respectively (Table 3). Additionally, CDPPB, DPFE and MPEP had significantly smaller maximum effects at human mGlu_5_ compared to rat mGlu_5_ (Table 3).

### 3.3. Assessment of allosteric modulation of glutamate and DHPG at human mGlu_5_ is affected by analysis approach

Next, modulatory activities for each allosteric modulator at human mGlu_5_ were investigated. Modulatory activity was only tested in iCa^2+^ mobilisation assays, as full intrinsic agonism and inverse agonism cannot be incorporated into the operational models of allosterism, precluding assessment using IP_1_ accumulation assays. Two approaches to analysing raw iCa^2+^ mobilisation fluorescent traces were undertaken; peak and area-under-the-curve (AUC). Peak values represent the maximum baseline-corrected fluorescence value recorded over the duration of the experiment under each ligand condition. However, taking a single peak value fails to capture the complexity of the full kinetic response to allosteric and orthosteric ligands. As such, AUC analysis was also used to define the total area of the ligand induced fluorescent response under each ligand condition. Concentration response curves derived from peak analysis of modulation of glutamate and DHPG are presented in Figures 3 and 4, with concentration-response curves derived from AUC analysis for glutamate modulation and DHPG modulation presented in Figures 5 and 6. All data were fitted to an operational model of allosterism to derive the functional affinity (pK_B_) and cooperativity (log_β_) estimates for each ligand for peak and AUC analyses, which are presented in Table 4 and Figure 5.

**Figure 3.**
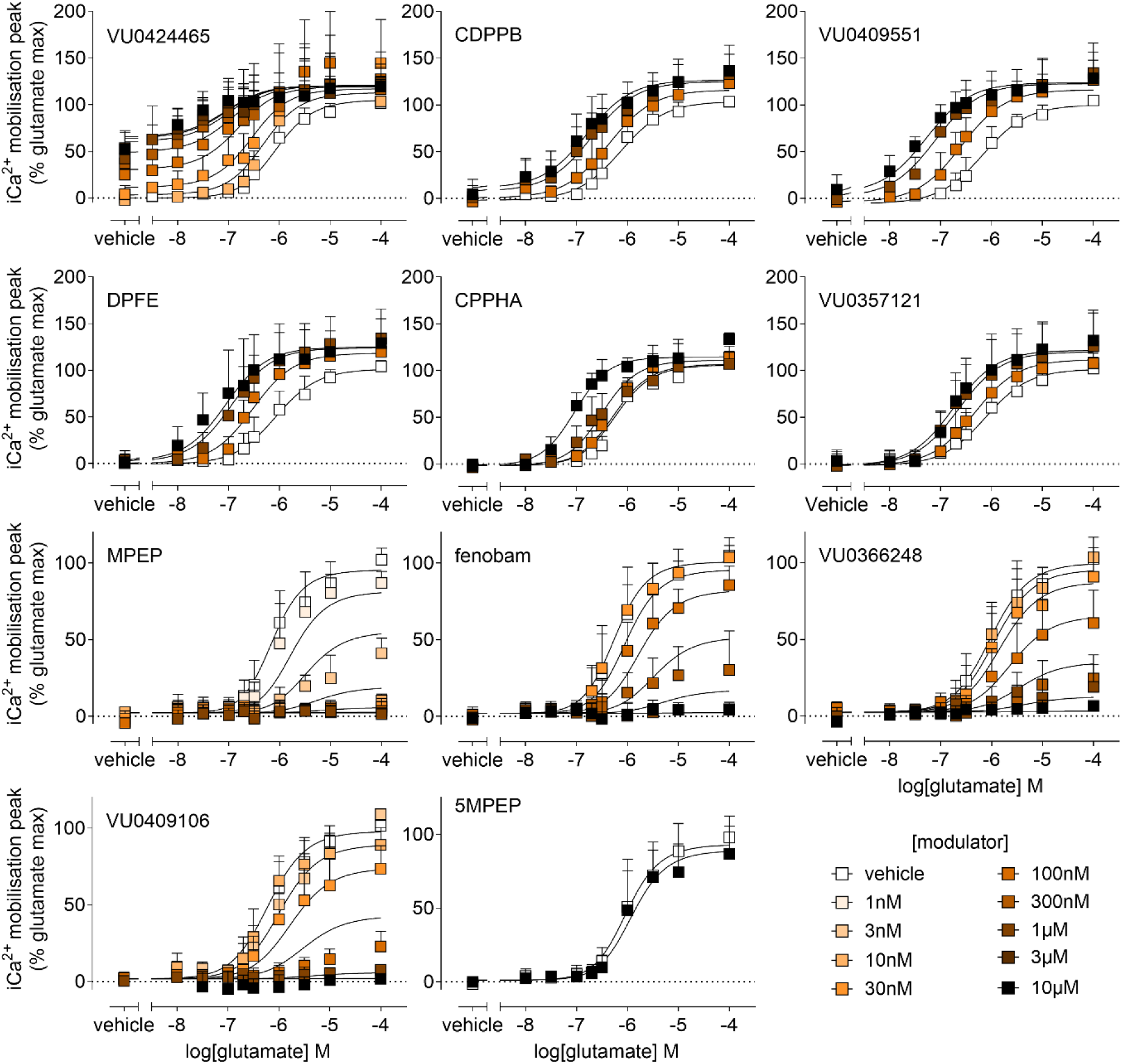
Peak analysis of allosteric modulation of glutamate-mediated iCa^2+^ mobilisation in HEK293A hmGlu_5_ cells. Glutamate concentration response curves were generated in the presence or absence of indicated concentrations of allosteric modulators, as denoted within each panel. All modulators were added 1 min prior to glutamate, with the exception of VU0424465, which was co-added with glutamate to minimise allosteric ligand-induced acute desensitisation. Data sets were globally fitted to a complete operational model of allosterism to estimate affinity and cooperativity. Curves represent the best fit of the data. Data are expressed as mean + S.D. of at least three experiments performed in duplicate. The number of independent experiments performed in duplicate for each ligand are listed in Table 3.

**Figure 4.**
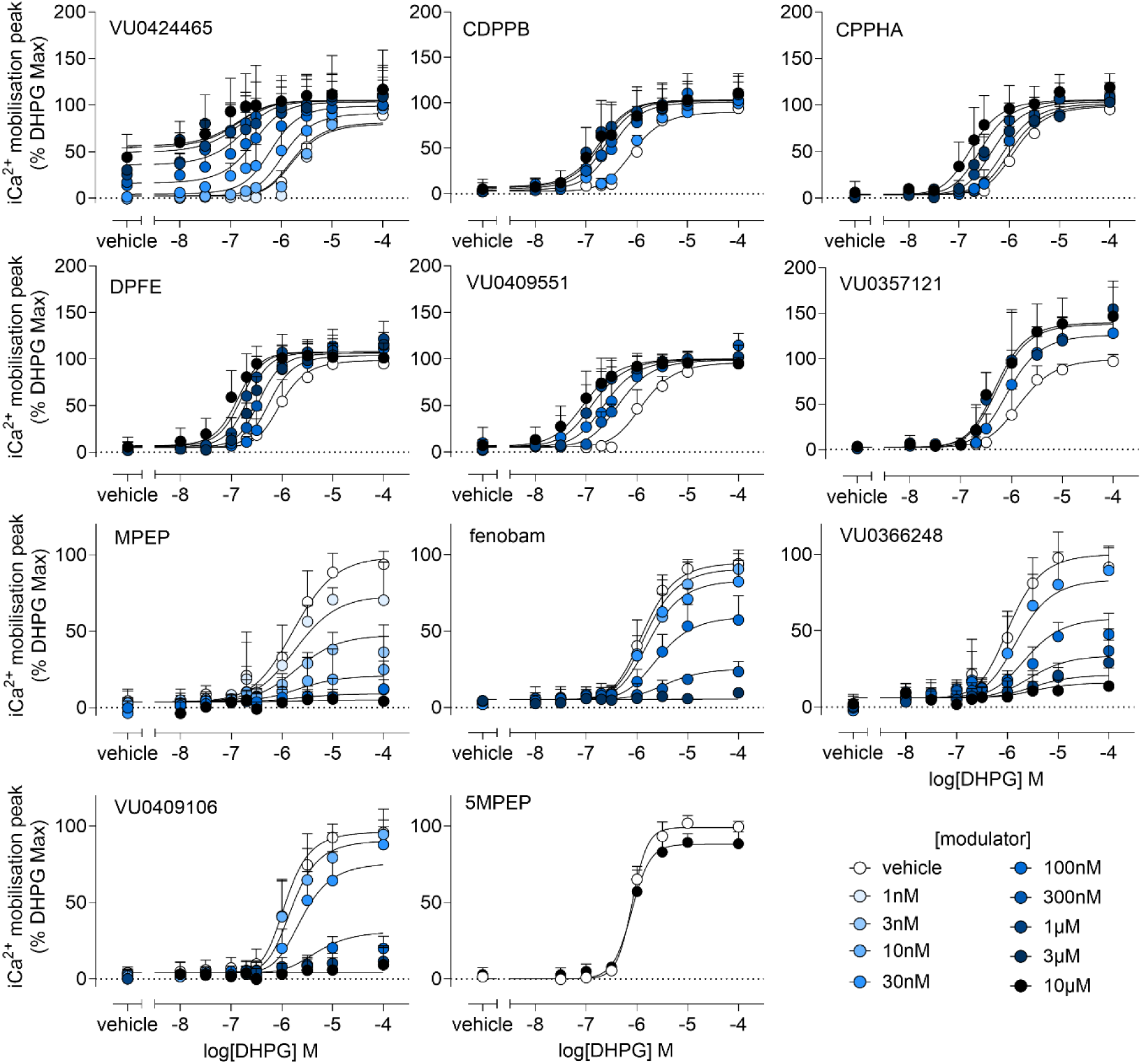
Peak analysis of allosteric modulation of DHPG-mediated iCa^2+^ mobilisation in HEK293A-hmGlu_5_ cells. DHPG concentration response curves were generated in the presence or absence of indicated concentrations of allosteric modulators, as denoted within each panel. All modulators were added 1 min prior to DHPG, with the exception of VU0424465, which was co-added with DHPG to minimise allosteric ligand-induced acute desensitisation. Data sets were globally fitted to a complete operational model of allosterism to estimate affinity and cooperativity. Curves represent the best fit of the data. Data are expressed as mean + S.D. of at least three experiments performed in duplicate. The number of independent experiments performed in duplicate for each ligand are listed in Table 3.

**Figure 5.**
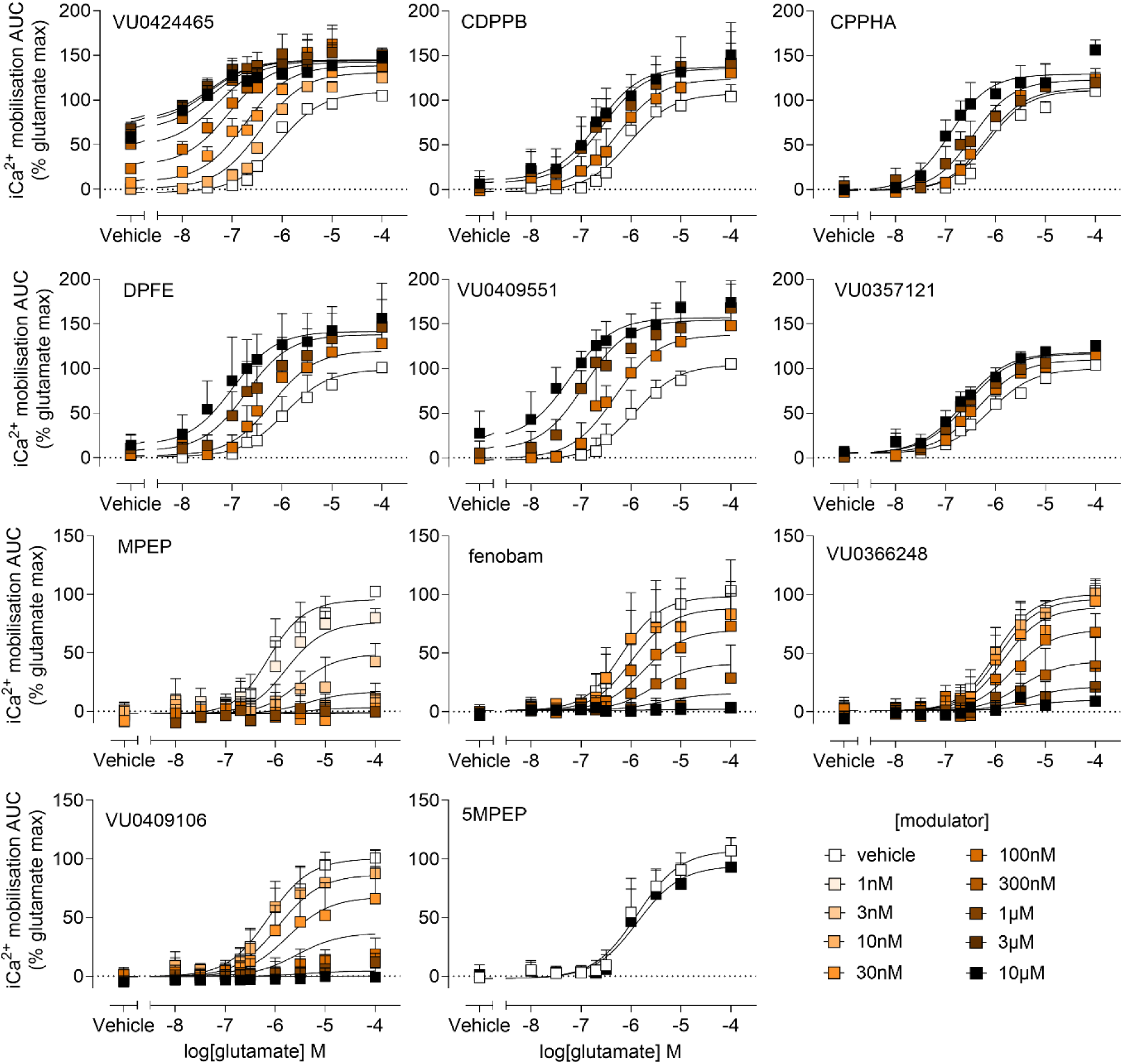
AUC analysis of allosteric modulation of glutamate-mediated iCa^2+^ mobilisation in HEK293A-hmGlu_5_ cells. Glutamate concentration response curves were generated in the presence or absence of indicated concentrations of allosteric modulators, as denoted within each panel. All modulators were added 1 min prior to glutamate, with the exception of VU0424465, which was co-added with glutamate to minimise allosteric ligand-induced acute desensitisation. Data sets were globally fitted to a complete operational model of allosterism to estimate affinity and cooperativity. Curves represent the best fit of the data. Data are expressed as mean + S.D. of at least three experiments performed in duplicate. The number of independent experiments performed in duplicate for each ligand are listed in Table 3.

**Figure 6.**
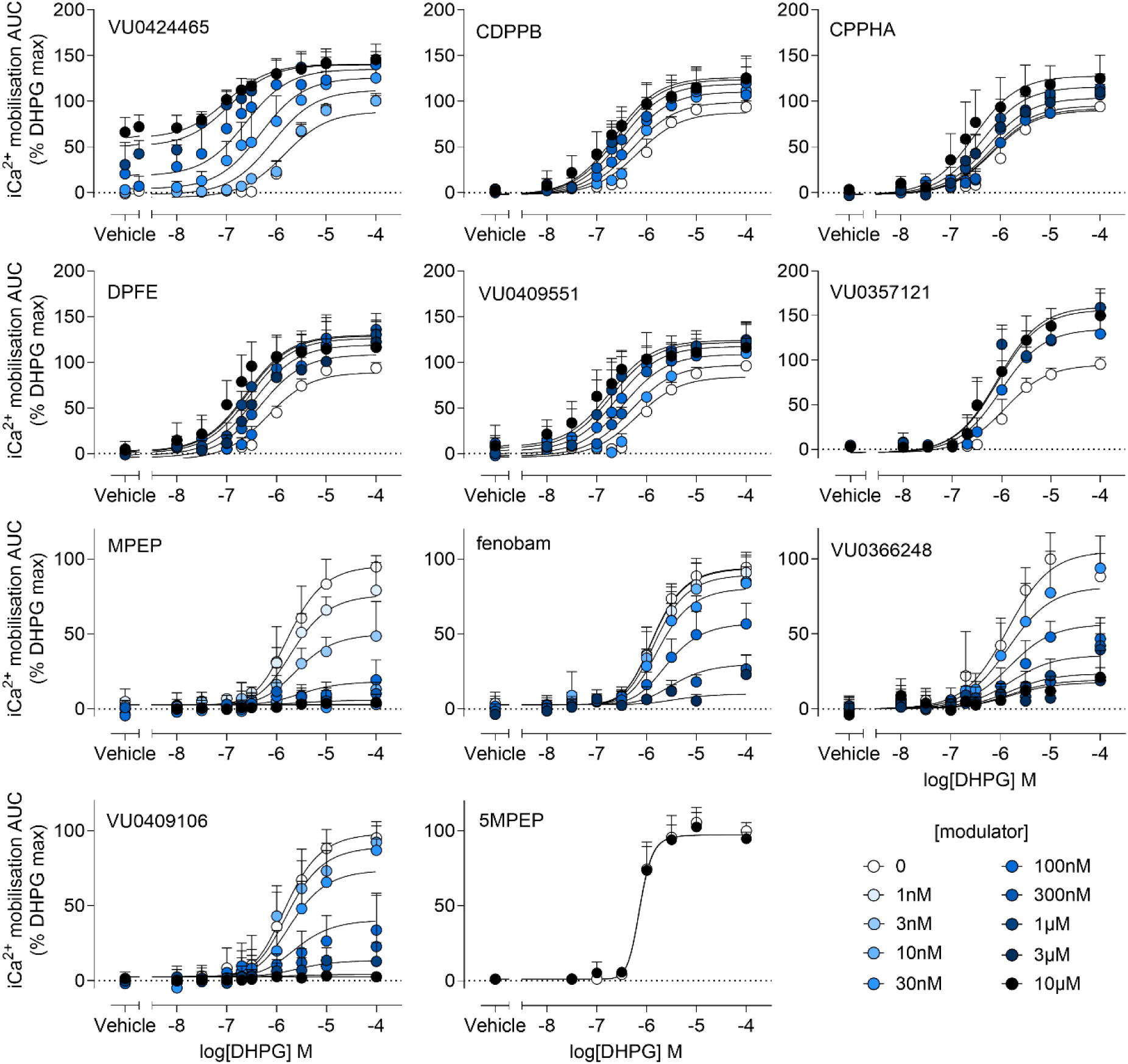
AUC analysis of allosteric modulation of DHPG-mediated iCa^2+^ mobilisation in HEK293A-hmGlu_5_ cells. DHPG concentration response curves were generated in the presence or absence of indicated concentrations of allosteric modulators, as denoted within each panel. All modulators were added 1 min prior to DHPG, with the exception of VU0424465, which was co-added with DHPG to minimise allosteric ligand-induced acute desensitisation. Data sets were globally fitted to a complete operational model of allosterism to estimate affinity and cooperativity. Curves represent the best fit of the data. Data are expressed as mean + S.D. of at least three experiments performed in duplicate. The number of independent experiments performed in duplicate for each ligand are listed in Table 3.

**Table 4.**
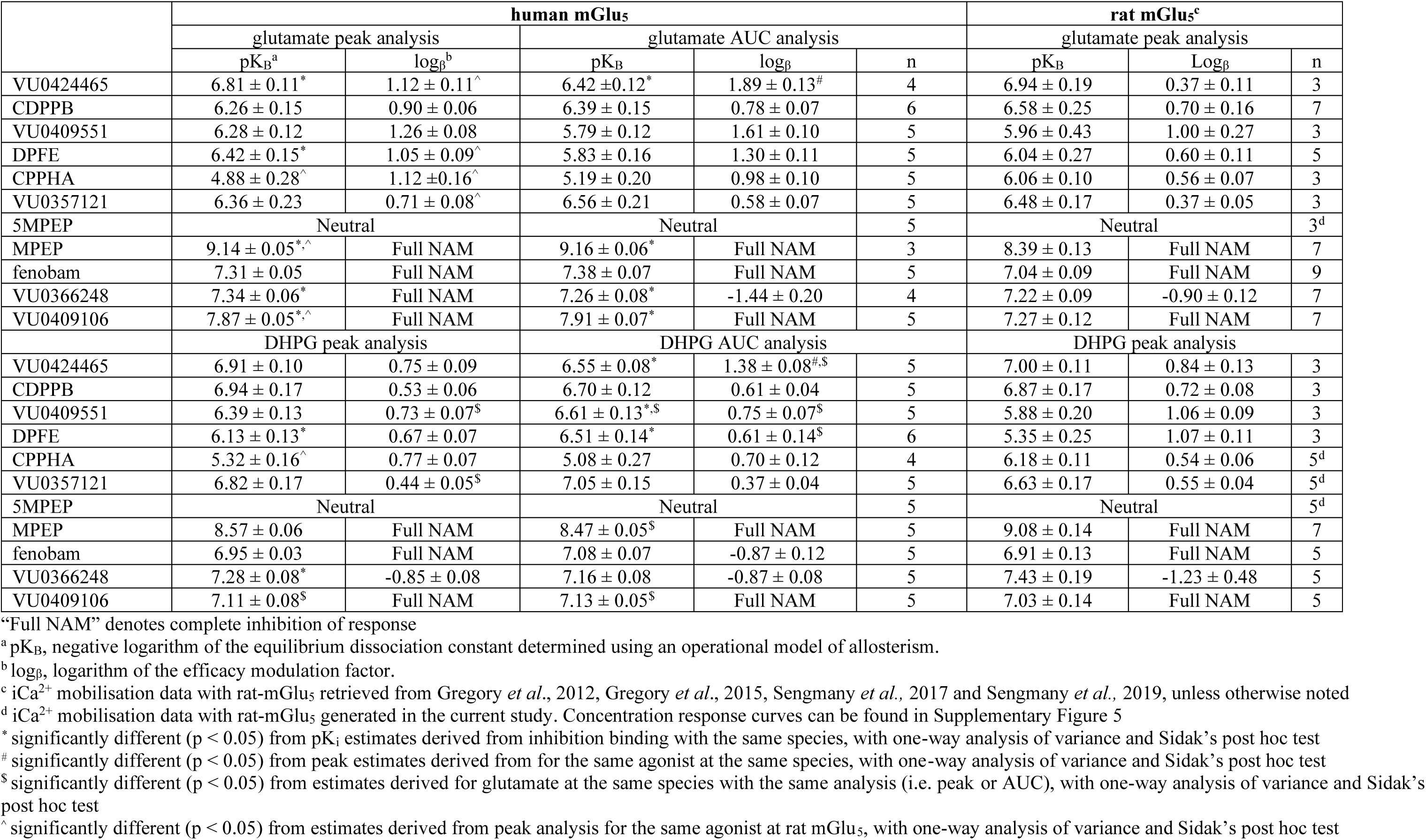
Comparison of affinity and cooperativity for allosteric modulation of orthosteric agonist-mediated iCa^2+^ mobilisation in HEK293A-hmGlu_5_ cells. Data represent mean ±SEM of the indicated number of independent experiments performed in duplicate.

Peak analysis revealed all PAMs induced a leftward shift in both glutamate and DHPG concentration-response curves for intracellular Ca^2+^ mobilisation (Figures 3 & 4). Additionally, all PAMs, with the exception of CPPHA, increased the maximum response to glutamate by 20-30%, and VU0357121 increased the maximum response to DHPG by 50%. NAMs inhibited glutamate and DHPG-stimulated iCa^2+^ mobilisation in a concentration-dependent manner (Figure 3 & 4). VU0366248, a partial NAM at rat mGlu_5_, completely inhibited glutamate-induced iCa^2+^ mobilisation, while acting as a partial NAM with high negative cooperativity for DHPG-induced iCa^2+^ mobilisation (Figure 3 & 4).

AUC analysis revealed mostly similar modulatory activity when compared to peak analysis, with some exceptions. PAM enhancement of the maximum glutamate effect was increased to ∼50% for all PAMs except VU0357121, and an enhancement of the DHPG maximum response was evident for all PAMs by ∼20-50% (Figures 5 and 6). VU0366248 acted as a partial NAM with high negative cooperativity for both glutamate- and DHPG-induced iCa^2+^ mobilization based on AUC analysis. No significant differences between analytical conditions were evident at the level of affinity for any modulator tested (Table 4). At the level of cooperativity, significant differences in logβ estimates were evident for VU0424465, with 4.3-fold and 6-fold higher logβ estimates for modulation of DHPG and glutamate, respectively (Table 4).

5MPEP has been reported as a NAL for glutamate-induced iCa^2+^ mobilisation at rat mGlu_5_ (Rodriguez *et al*., 2005). To confirm this profile, here 5MPEP was tested for modulation of glutamate and DHPG-induced iCa^2+^ mobilisation at rat mGlu_5_. 5MPEP was confirmed as a NAL for DHPG at rat mGlu_5_ (Figure 7). Similarly, at human mGlu_5,_ 5MPEP acted as a NAL for modulation of glutamate, and only marginally reduced the maximum response to DHPG at 10µM by ∼10% (Figures 3-6).

**Figure 7.**
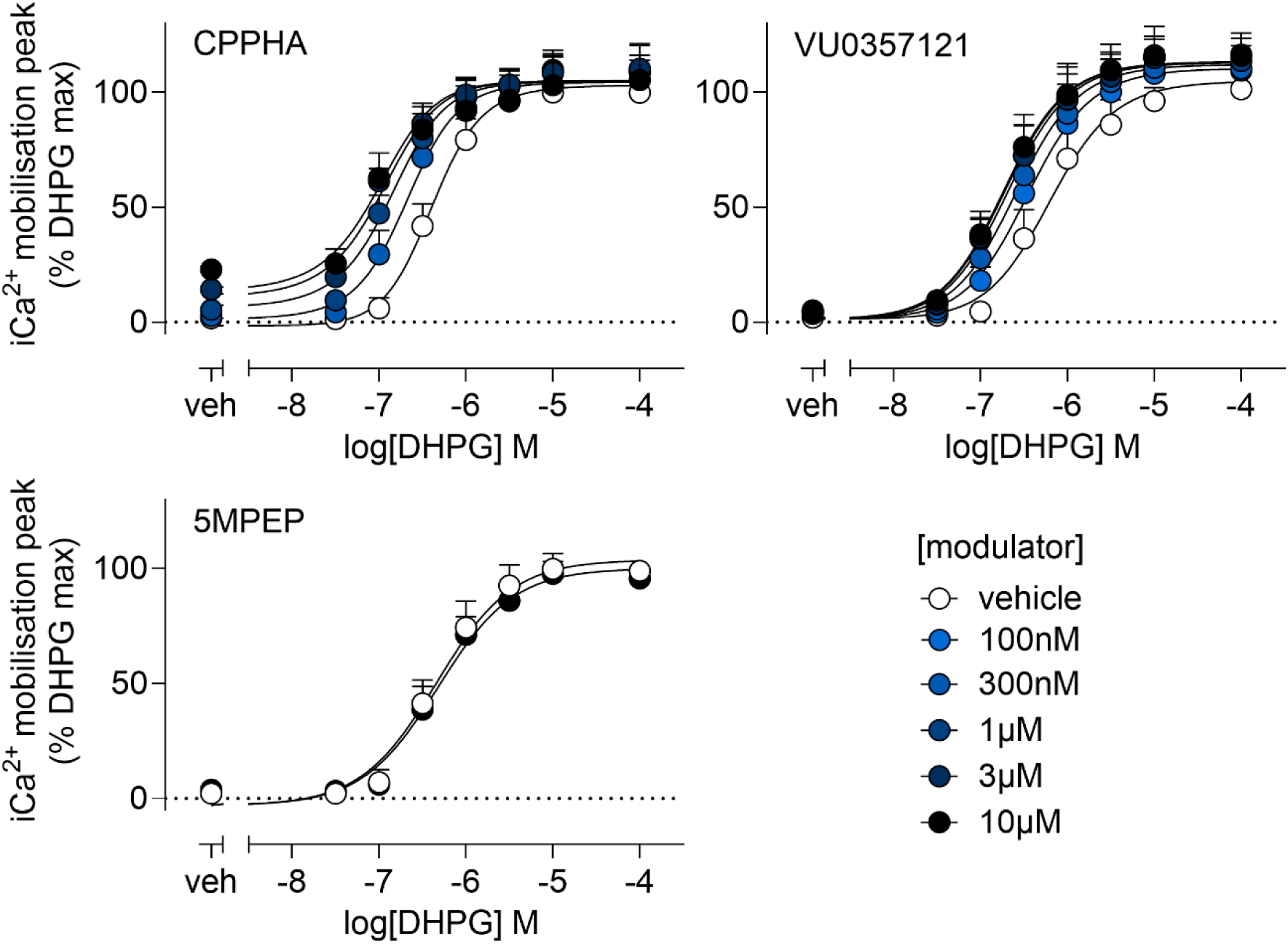
Peak analysis of allosteric modulation of DHPG-mediated iCa^2+^ mobilisation in HEK293A-rmGlu_5_ cells. DHPG concentration response curves were generated in the presence or absence of indicated concentrations of allosteric modulators, as denoted within each panel, added 1 min prior to DHPG. Data sets were globally fitted to a complete operational model of allosterism to estimate affinity and cooperativity. Curves represent the best fit of the data. Data are expressed as mean + S.D. of at least three experiments performed in duplicate. The number of independent experiments performed in duplicate for each ligand are listed in Table 3.

Functional pK_B_ values were next compared to pK_i_ estimates derived from binding experiments, to determine the correlation between functional and binding affinity. Functional pK_B_ estimates for VU0424465 were significantly lower than binding pK_i_ estimates when comparing both peak (4.5-fold) and AUC analysis (10-fold) of glutamate modulation, and AUC analysis of DHPG modulation (8-fold; Figure 8A). Functional affinity values for DPFE were significantly higher than binding affinity estimates when comparing peak and AUC analysis of DHPG modulation (10- and 24-fold, respectively), and peak analysis of glutamate modulation (20-fold; Figure 5A). Interestingly, functional affinity estimates were significantly higher for both analyses of glutamate modulation for both MPEP (5.5-fold) and VU0409106 (10-fold), but not for DHPG modulation, with VU0366248 displaying increased functional affinity for modulation of both glutamate (5-fold) and DHPG (3-fold) relative to binding affinity (Figure 8A). Correlation analysis revealed pK_B_ estimates derived from peak analysis of DHPG modulation was most highly correlated with binding derived pK_i_ estimates (slope = 0.72 ± 0.15; r^2^ = 0.72), followed by glutamate AUC analysis (slope = 0.96 ± 0.23; r^2^ = 0.67), glutamate peak analysis (slope = 0.92 ± 0.23; r^2^ = 0.65) and DHPG AUC analysis (slope = 0.62 ± 020; r^2^ = 0.51) (Figure 5B).

**Figure 8.**
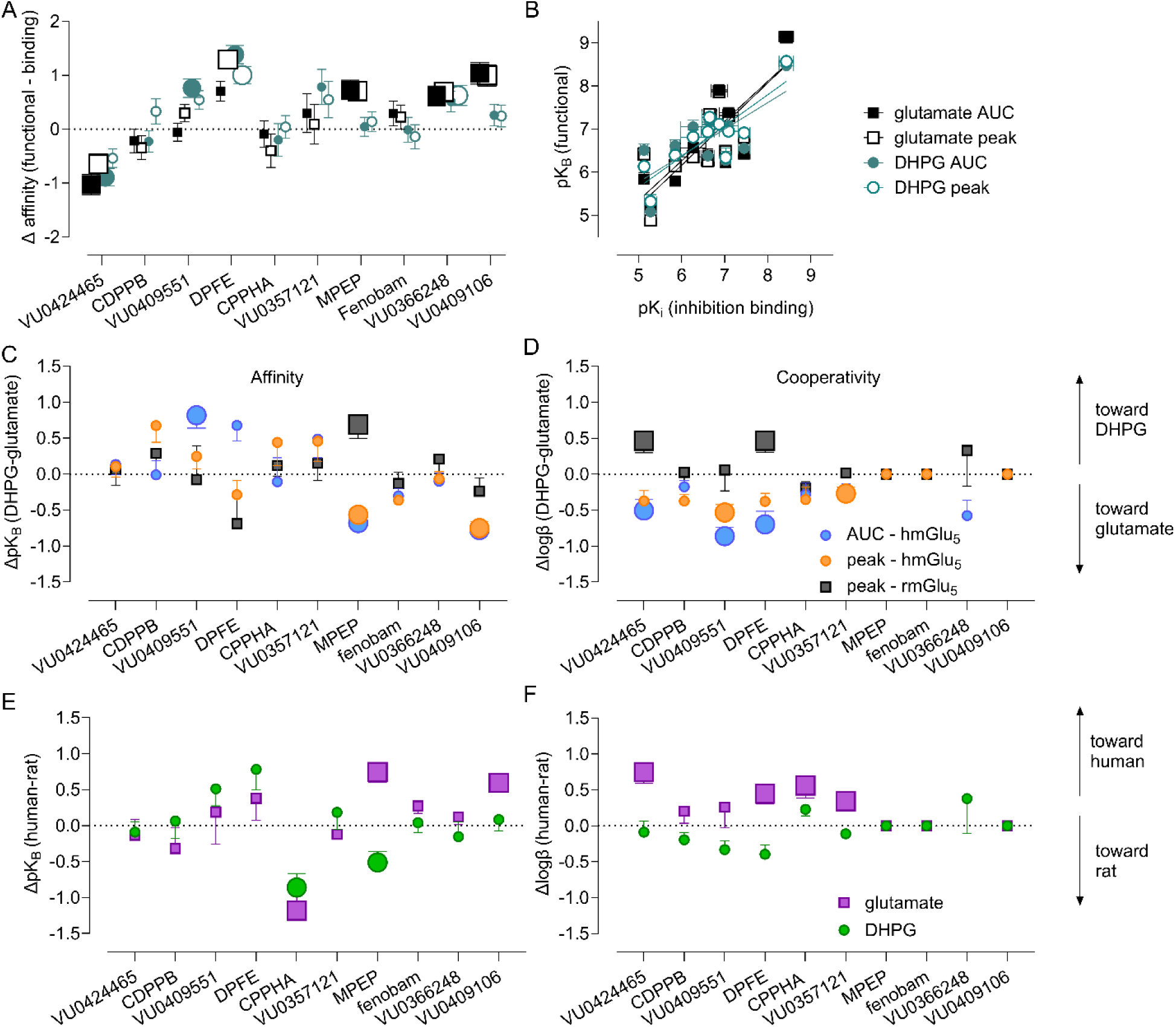
Comparison of affinity and cooperativity for mGlu_5_ allosteric modulators derived from binding and iCa^2+^ mobilisation assays, using both area under curve (AUC) and peak analysis. (**A**) Δaffinity estimates for each ligand derived from binding and both glutamate and DHPG-mediated iCa^2+^ mobilisation assay. Large symbols denote statistically significant differences between functional and binding affinity estimates. (**B**) Correlation plot for functional pK_B_ estimates relative to inhibition binding derived pK_i_ values for mGlu_5_ allosteric modulators. (**C & D**) Probe dependence is evident for affinity (C) and cooperativity (D) estimates for each ligand derived from peak and AUC analysis of indicated orthosteric ligand-based iCa^2+^ mobilisation assays in HEK293A-hmGlu_5_ cells and peak analysis of orthosteric ligand-based iCa^2+^ mobilisation assays in HEK293A-rmGlu_5_. Large symbols denote statistically significant differences between estimates derived from modulation of DHPG and glutamate. (**E & F**) Evaluation of species differences for affinity (E) and cooperativity (F) estimates for each ligand derived from peak analysis of indicated orthosteric ligand-based iCa^2+^ mobilisation assays in HEK293A-hmGlu_5_ and HEK293A-rmGlu_5_. Large symbols denote statistically significant differences between estimates derived from modulation of human mGlu_5_ and rat mGlu_5_. Affinity and cooperativity for rat mGlu_5_ were retrieved from Gregory *et al.,* 2012, Gregory *et al.,* 2015, Sengmany *et al.,* 2017 and Sengmany *et al.,* 2019 or generated on the current study (see Table 3).

### 3.4. mGlu_5_ allosteric modulators display probe- and species-dependent pharmacology

Probe dependence is a phenomenon in which allosteric modulators display different pharmacological properties depending on the orthosteric agonist present, and is evident for mGlu_5_ PAMs and NAMs at the level of both affinity and cooperativity at rat mGlu_5_ (Sengmany *et al*., 2017, 2019). At the level of affinity, the only PAM to display probe dependence at human mGlu_5_ was VU0409551, with AUC analysis revealing a 7-fold higher affinity estimate for modulation of DHPG compared to glutamate (Figure 8C). At the level of affinity, MPEP and VU0409106 showed probe dependence, with 4-5-fold and 5.5-fold higher affinity estimates for modulation of DHPG relative to glutamate, respectively (Figure 8C). At the level of cooperativity, AUC analysis revealed significantly higher logβ estimates for modulation of glutamate, relative to DHPG, for VU0424465 (3.5-fold), VU0409551 (7-fold) and DPFE (5-fold) (Figure 8D). Peak analysis revealed significantly higher logβ estimates for modulation of glutamate, relative to DHPG, for VU0409551 (3-fold) and VU0357121 (2-fold) (Figure 8D).

When compared to estimates derived from peak analysis of modulation of glutamate and DHPG-induced iCa^2+^ mobilisation at rat mGlu_5,_ affinity and cooperativity of all compounds at human mGlu_5_ were similar to those previously determined, with some species-specific differences (Table 4, Figure 8E & F). Affinity estimates were mostly similar for PAMs, with the exception of CPPHA, which had a 15-fold lower affinity for modulation of glutamate and 7-fold lower affinity for modulation of DHPG for human mGlu_5_ compared to rat mGlu_5_ (Figure 8E). Conversely, MPEP and VU0409106 had higher functional affinity for the human receptor when modulating glutamate, by 5.5-fold and 4-fold, respectively (Figure 8E). MPEP also had 3-fold lower functional affinity for the human receptor when modulating DHPG. For MPEP, these differences resulted in probe dependence switching direction between human and rat mGlu_5_, with probe dependence at the toward DHPG for rat mGlu_5_ and glutamate for human mGlu_5_ (Figure 8C). VU0409106 also gained probe dependence toward glutamate at human mGlu_5_, which was not evident at rat mGlu_5._ When comparing cooperativity, logβ estimates for modulation of glutamate were different for most PAMs, being significantly higher at the human mGlu_5_ for VU0424465 (5.5-fold), DPFE (3-fold), CPPHA (3.5-fold) and VU0357121 (2-fold).

### 3.5. mGlu_5_ positive allosteric modulators differentially affect orthosteric agonist kinetic profiles

The difference in pharmacological parameters between modulation of different agonists and between AUC and peak analyses may arise from allosteric modulators differentially changing the kinetics of agonist induced iCa^2+^ responses. To investigate this, kinetic analyses were carried out on time traces for glutamate- and DHPG-induced iCa^2+^ responses in the absence and presence of allosteric modulators (Figures 9 and 10). Limitations with kinetic models meant responses with a shallow rise phase were unable to be fit, such as those for low concentrations of agonist or agonist in the presence of NAMs, therefore only the highest concentration of each agonist was analysed in the absence and presence of PAMs. To ensure equivalent conditions were compared across different PAMs, agonist responses were compared in the presence of an approximate K_B_/K_i_ concentration (∼50% occupancy; based on PAM binding affinity estimates from Table 2) and a maximum concentration (at least 10x K_B_/K_i_, >90% occupancy; 10µM for each PAM). For CPPHA and DPFE, only K_B_ values were analysed as these approached 10µM. Three kinetic parameters were estimated (Figure 11A); initial rate (steepness of the inflection of the rise phase), K_1_ (rate constant of the rise phase) and K_2_ (rate constant of the decay phase). To account for differences in kinetics between independent experiments, different PAM concentrations were compared to vehicle controls from within the same experiment. Kinetic parameter estimates are displayed in Table 5 and normalised (Δ) estimates in Figure 11. Some ligand combinations could not be fit and are noted in the Tables. At respective K_B_ concentrations, CPPHA and CDPPB significantly enhanced glutamate initial rate, without affecting DHPG, while the opposite was true for VU0357121 (Figure 11B). VU0409551 significantly enhanced the initial rate of both agonists, while DPFE and VU0424465 had no effect on initial rate at K_B_ concentrations. At 10µM, CDPPB, VU0409551 and VU0424465 significantly enhanced the initial rate of both agonists, whereas VU0357121 only affected the initial rate of DHPG (Figure 11B). A significant reduction in K_1_ was observed for DHPG in the presence of K_B_ concentrations of CDPPB, DPFE and both concentrations of VU0424465, whereas both concentrations of VU0409551 significantly increased DHPG K_1_ (Figure 11C). The only differences in glutamate K_1_ were a significant increase in the presence of CDPPB, and a significant decrease in the presence of DPFE (Figure 11C). CPPHA, CDPPB and DPFE had no effect on K_2_ estimates for either agonist (Figure 11D). At K_B_ concentrations, VU0409551 significantly decreased both DHPG and glutamate K_2_ estimates, whereas VU0424465 caused a significant decrease and a significant increase in K_2_ estimates for glutamate and DHPG, respectively (Figure 11D). Similarly, 10µM VU0409551 significantly decreased, where 10µM VU0424465 significantly increased, K_2_ estimates for DHPG (Figure 11D).

**Figure 9.**
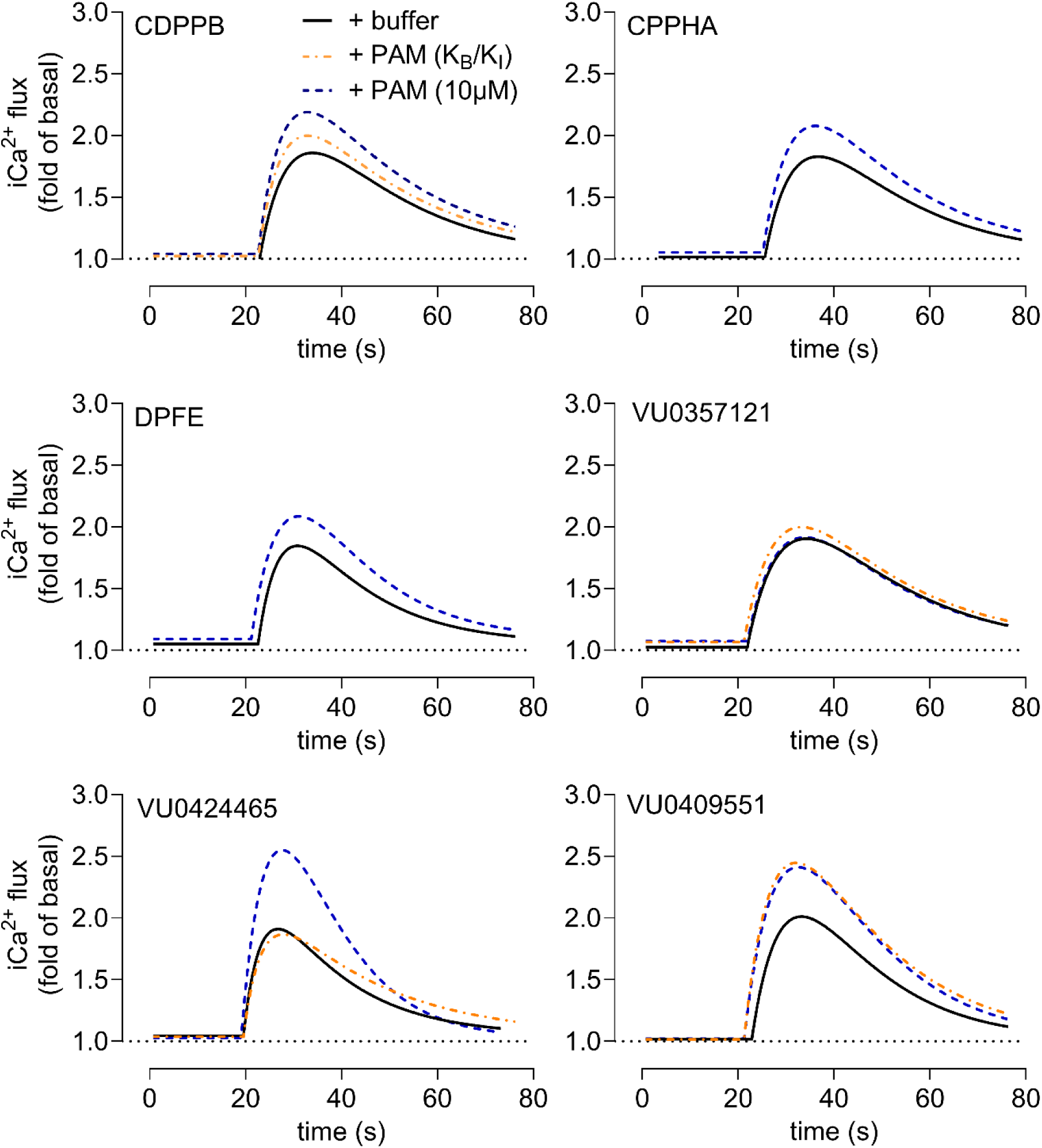
Kinetic traces (time vs RFU fold change) for glutamate-induced iCa^2+^mobilisation in the absence and presence of different mGlu_5_ PAMs. Time vs. ΔRFU (fold over baseline) data for iCa^2+^ mobilisation induced by 100µM glutamate were plotted and fitted to the kinetic “rise-then-fall to baseline” kinetic model (equation 7). Glutamate in the presence of vehicle (solid black lines) were then compared to the presence of different mGlu_5_ PAMs at concentrations approaching their affinity values (determined from binding assays; orange dashed lines) or at 10µM (high occupancy; blue dashed lines). See Table 5 for derived kinetic parameters. Data represent best-fit to iCa^2+^ mobilisation time traces from 4-6 independent experiments. Vehicle curves are matched controls from within independent experiment for each PAM.

**Figure 10.**
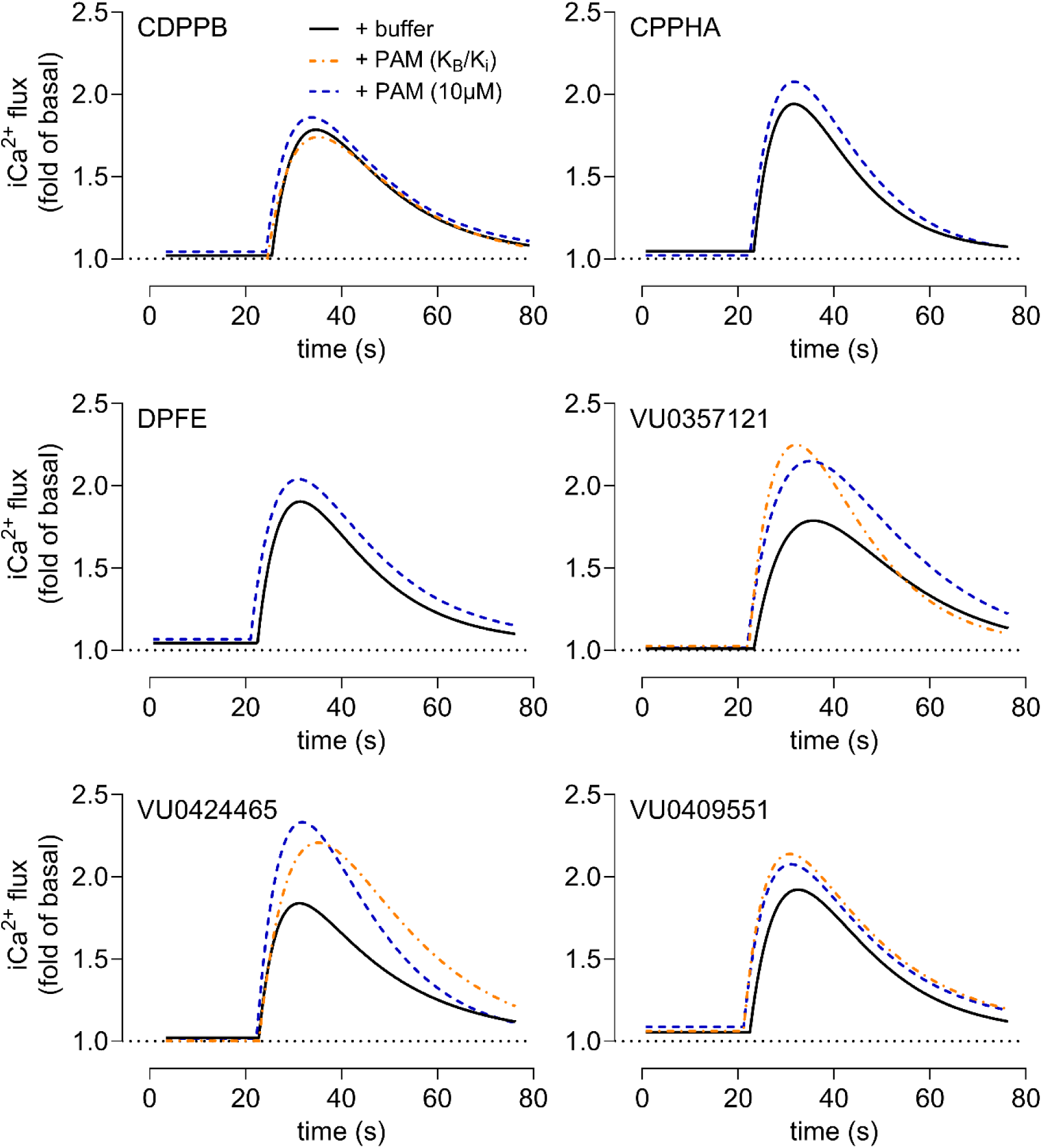
Kinetic traces (time vs RFU fold change) for DHPG-induced iCa^2+^mobilisation in the absence and presence of different mGlu_5_ PAMs. Time vs. ΔRFU (fold over baseline) data for iCa^2+^ mobilisation induced by 30µM DHPG were plotted and fitted to the kinetic “rise-then-fall to baseline” kinetic model (equation 7). DHPG in the presence of vehicle (solid black lines) were then compared to the presence of different mGlu_5_ PAMs at concentrations approaching their affinity values (determined from binding assays; orange dashed lines) or at 10µM (high occupancy; blue dashed lines). See Table 5 for derived kinetic parameters. Data represent best-fit to iCa^2+^ mobilisation time traces from 4-6 independent experiments. Vehicle curves are matched controls from within independent experiment for each PAM.

**Figure 11.**
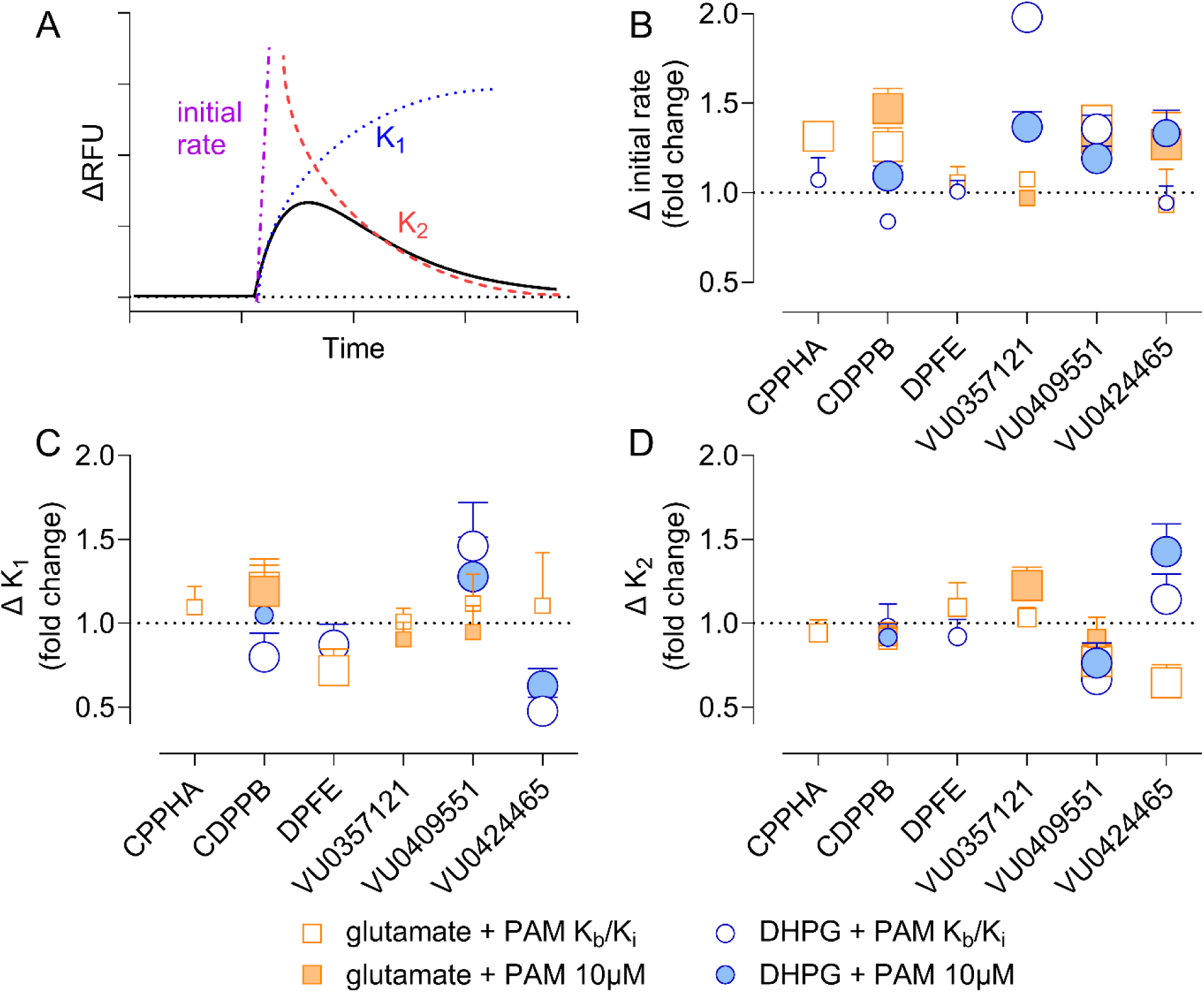
Derivation and comparison of kinetic parameters for glutamate and DHPG-induced iCa^2+^ mobilisation in the absence and presence of positive allosteric modulators in HEK293A-hmGlu_5_ cells. (**A**) schematic of best-fit kinetic curves fit to iCa^2+^ mobilisation data. Time vs. ΔRFU (fold over baseline) was plotted and fitted to the kinetic “rise-then-fall to baseline” kinetic model (equation 7). Parameters initial rate (slope of initial response) and K_1_ (rate constant of the rise phase) and K_2_ (rate constant of the fall phase) were derived and compared between orthosteric agonists in the presence of vehicle or differing concentrations of PAMs (see Table 5). Initial rate, K_1_ and K_2_ estimates from kinetic curves in the presence of PAMs were normalised to estimates in the presence of vehicle to derive Δ values, plotted in panels B-D. (**B**) PAMs increase the initial rate of orthosteric ligand induced iCa^2+^ signalling in a PAM and agonist dependent manner. (**C**) PAMs change rise-phase rate constant for DHPG in a ligand-dependent manner, with little effect on glutamate K_1_ estimates (**D**) Only VU0357121, VU0409551 and VU0424465 significantly affect the rate constant of the fall-phase for either agonist. Large symbols denote statistically significant differences between estimates derived in the absence and presence of indicated concentrations of PAMs.

**Table 5:** Kinetic parameters for orthosteric agonist-induced iCa^2+^ mobilisation in the absence and presence of mGlu_5_ positive allosteric modulators. Data represent mean ±SEM of the indicated number of independent experiments performed in duplicate.

| | agonist (100$\mu\text{M}$) + vehicle | | | agonist (100$\mu\text{M}$) + PAM ($K_i/K_B$)<sup>a</sup> | | | agonist (100) + PAM (10$\mu\text{M}$) | | | |
| --- | --- | --- | --- | --- | --- | --- | --- | --- | --- | --- |
| | Initial rate ($\text{s}^{-1}$) | K1 | K2 | Initial rate ($\text{s}^{-1}$) | K1 | K2 | Initial rate ($\text{s}^{-1}$) | K1 | K2 | n |
| glutamate |  |  |  |  |  |  |  |  |  |  |
| VU0424465 | 0.35±0.05 | 0.26±0.05 | 0.06±0.01 | 0.33±0.05 | 0.28±0.05 | 0.04±0.004* | 0.45±0.01* | n.d. | n.d. | 4 |
| CDPPB | 0.22±0.01 | 0.15±0.02 | 0.05±0.004 | 0.28±0.02* | 0.19±0.02* | 0.04±0.002 | 0.33±0.02* | 0.18±0.02* | 0.04±0.002 | 6 |
| VU0409551 | 0.27±0.01 | 0.13±0.02 | 0.07±0.01 | 0.37±0.01* | 0.14±0.01 | 0.05±0.003* | 0.35±0.01* | 0.12±0.01 | 0.06±0.005 | 5 |
| DPFE | 0.27±0.02 | 0.19±0.02 | 0.07±0.01 | 0.28±0.01 | 0.14±0.02* | 0.07±0.01 | n.a. | n.a. | n.a. | 5 |
| CPPHA | 0.20±0.01 | 0.14±0.01 | 0.05±0.003 | 0.27±0.01* | 0.15±0.02 | 0.05±0.003 | n.a. | n.a. | n.a. | 5 |
| VU0357121 | 0.20±0.01 | 0.12±0.01 | 0.05±0.002 | 0.22±0.004 | 0.12±0.01 | 0.05±0.001 | 0.20±0.01 | 0.11±0.01 | 0.06±0.005* | 5 |
| DHPG |  |  |  |  |  |  |  |  |  |  |
| VU0424465 | 0.29±0.03 | 0.22±0.03 | 0.05±0.004 | 0.27±0.01 | 0.11±0.01* | 0.06±0.005* | 0.38±0.01* | 0.14±0.01* | 0.08±0.01* | 5 |
| CDPPB | 0.23±0.02 | 0.16±0.03 | 0.07±0.01 | 0.20±0.01 | 0.13±0.02* | 0.07±0.01 | 0.26±0.01* | 0.17±0.01 | 0.06±0.003 | 5 |
| VU0409551 | 0.24±0.01 | 0.12±0.02 | 0.08±0.01 | 0.32±0.01* | 0.18±0.01* | 0.05±0.003* | 0.28±0.02* | 0.16±0.01* | 0.06±0.004* | 5 |
| DPFE | 0.27±0.01 | 0.16±0.02 | 0.07±0.01 | 0.27±0.01 | 0.14±0.01* | 0.07±0.004 | n.a. | n.a. | n.a. | 6 |
| CPPHA | 0.29±0.03 | n.d. | n.d. | 0.31±0.02 | n.d. | n.d. | n.a. | n.a. | n.a. | 4 |
| VU0357121 | 0.17±0.01 | n.d. | n.d. | 0.34±0.01* | n.d. | n.d. | 0.23±0.01* | n.d. | n.d. | 5 |

<sup>a</sup> agonist in the presence of PAMs at concentrations approximating affinity values derived from binding assays; CPPHA = 10 $\mu\text{M}$ , CDPPB = 100nM, DPFE = 10 $\mu\text{M}$ , VU0409551 = 1 $\mu\text{M}$ , VU0424465 = 30nM, VU0357121 = 1 $\mu\text{M}$
n.a. not applicable as affinity values are approximately 10 $\mu\text{M}$
n.d. not determined as kinetic curves could not be fit
\* $p > 0.05$ compared to estimates for agonists in the presence of vehicle from matched experimental datasets, determined using a sum-of-squares F-test

## 4. Discussion

Despite preclinical success in rodent models of multiple CNS disorders, development of mGlu_5_ targeted drugs has stalled, either due to lack of efficacy or concerns over adverse effects at both the preclinical and clinical stages (Bridges *et al*., 2013; Parmentier-Batteur *et al*., 2014; Gould *et al*., 2016; Barnes *et al*., 2018; Budgett *et al*., 2022; Luessen and Conn, 2022). Translational failures have been suggested to arise from drug discovery programs categorising novel compounds based on a single functional assay (i.e. iCa^2+^ mobilisation), over reliance on use of rodent receptor and the subsequent scarcity of preclinical data at human mGlu_5_ (Sengmany *et al*., 2017, 2019; Hellyer *et al*., 2019; Arsova *et al*., 2020, 2021). Such limitations may ignore the complexity of allosteric modulator pharmacology and hamper accurate linking of *in vitro* and *in vivo* profiles. Here we performed quantitative pharmacological characterisation of eleven allosteric modulators in HEK293A cells stably expressing human mGlu_5_ using radioligand inhibition binding, iCa^2+^ mobilisation and IP_1_ accumulation assays. The eleven modulators chosen were previously categorised as ago-PAMs, second-site PAMs, NAMs and NALs for rat mGlu_5._ We reveal species-dependent differences in binding and pharmacology for structurally and pharmacology diverse ligands. Many PAMs lacked agonism for iCa^2+^ mobilisation at human mGlu_5_ and were significantly less potent in IP_1_ accumulation when compared to rat mGlu_5._ Modulatory studies revealed probe dependence was reversed for select ligands when comparing between species. Overall, our findings reiterate the need to characterise mGlu_5_ allosteric modulator pharmacology at the translationally relevant human mGlu_5_, to enrich our understanding of pharmacological complexity and to provide a deeper insight into species differences that may impact translatability from preclinical to clinical settings.

Estimates of mGlu_5_ allosteric modulator affinity were mostly conserved across species, with significant differences between rat and human only evident for VU0424465 and VU0409106. Despite binding to the same common “MPEP” binding site on mGlu_5_ as the radioligand, many modulators did not fully displace [^3^H]methoxy-PEPy at human mGlu_5._ Incomplete radioligand displacement commonly indicates non-competitive interactions between the radioligand and competing ligand (Pagano *et al*., 2000; Flanagan, 2016). For CPPHA and VU0357121 this was expected, as these PAMs bind to as-yet unidentified sites distinct from the common binding site (Chen *et al*., 2008; Hammond *et al*., 2010; Gregory *et al*., 2012). VU0409106 and VU0366248 have also previously been reported to incompletely inhibit whole cell [^3^H]methoxy-PEPy binding at rat mGlu_5_ and mouse cortical neurons (Sengmany *et al*., 2019). However, previous binding experiments at rat mGlu_5_ reported full inhibition of [^3^H]methoxy-PEPy binding by the PAM CDPPB (Gregory *et al*., 2012). The use of membrane preparations in previous studies may play a role in the different binding profiles, as previously suggested allosteric modulators may stabilise receptor conformations only found in intact cells due to interactions between mGlu_5_ and intracellular signalling and scaffolding partners (Sengmany *et al*., 2019). Alternatively, the presence of two site binding for MPEP in the current study, and for MPEP and fenobam at rat mGlu_5_ in previous studies, suggest ligands incompletely displacing [^3^H]methoxy-PEPy may have low affinity for this second site (Sengmany *et al*., 2019). Technical artifacts may also play a role, as non-specific binding is defined with MPEP, which may be competing for non-specific binding sites with modulators of the same/similar chemotypes. As mGlu_5_ is known to be highly expressed on intracellular membranes as well as the plasma membrane, and allosteric modulators with distinct structures likely have different membrane permeability, differential access to distinct receptor pools may play a role in both incomplete radioligand displacement and the presence of a potential second-site when measured using whole cell preparations (Jong *et al*., 2009, 2019; Vincent *et al*., 2016). It is also possible these effects may be influenced by differential binding across the mGlu_5_ dimer. Binding to one protomer may affect the affinity of the other protomer for allosteric ligands, or modulators may display divergent occupancy of binding sites within each protomer (e.g. two molecules per protomer). Despite these subtle differences, the data presented here indicates minimal species dependence for mGlu_5_ allosteric ligand binding.

Both DHPG and PAM agonists had higher efficacy and potency in IP_1_ accumulation assays compared to iCa^2+^ mobilisation assays at human mGlu_5_. NAMs also lacked inverse agonist activity for iCa^2+^ mobilisation but acted as inverse agonists for IP_1_ accumulation. These data support previous observations from both rat mGlu_5_ and both rat and human orthologs of the closely related mGlu_1_, in which orthosteric and allosteric ligands displayed higher agonist or inverse agonist activity in IP_1_ accumulation, relative to iCa^2+^ mobilisation, assays (Sengmany *et al*., 2017, 2020; Hellyer *et al*., 2019, 2020; Arsova *et al*., 2020, 2021; Muraleetharan *et al*., 2023). Such differences between assays have been suggested to be linked to constitutive mGlu_5_ activity, ligand binding kinetics and interplay between receptor activation and regulatory processes (Sengmany *et al*., 2017, 2019; Muraleetharan *et al*., 2023). Slow binding ligands and those that enhance/inhibit constitutive activity would be more likely to have activity in equilibrium (i.e. IP_1_ accumulation) versus transient (i.e. iCa^2^ mobilisation) assays, as ligand efficacy and receptor constitutive activity are likely proportional to the duration of ligand-receptor interactions and receptor-G protein interactions, respectively (Parmentier *et al*., 1998; Carroll *et al*., 2001; Tummino and Copeland, 2008; Klein Herenbrink *et al*., 2016; Lane *et al*., 2017). Although relative differences in potencies between assays was conserved between species, DHPG and the PAMs VU0424465, DPFE and CDPPB had significantly lower potency estimates for human mGlu_5_ in both iCa^2+^ mobilisation and IP_1_ accumulation assays compared to rat mGlu_5_. While potency differences for VU0424465 may be linked to lower binding affinity for human mGlu_5_, allosteric agonism has also been attributed to increased mGlu_5_ receptor reserve, and differences in receptor reserve between HEK293A-hmGlu_5_ and HEK293A-rmGlu_5_ cell lines may explain the observed lower potency for other PAMs (Noetzel et al., 2012). However, VU0409551 and all NAMs had similar potency estimates for both species in IP_1_ accumulation, indicating divergent effects among distinct chemotypes and highlighting the importance of testing multiple signalling outcomes across different species for mGlu_5_.

Modulatory activity of mGlu_5_ PAMs and NAMs were mostly conserved between species. For four of the six PAMs tested, cooperativity was higher for glutamate modulation at human mGlu_5_ relative to rat mGlu_5_. Additionally, apparent affinity was higher for modulation of glutamate at human mGlu_5_ for MPEP and VU0409106, relative to rat mGlu_5._ Such differences had subsequent effects on observed probe dependence. Surrogate agonists are often used in native cell preparations and investigating probe dependence is crucial to translating findings from recombinant systems to native preparations. Probe dependence is evident for both cooperativity and functional affinity estimates of mGlu_5_ allosteric modulators at rat mGlu_5_ (Sengmany *et al*., 2017, 2019; Hellyer *et al*., 2020; Arsova *et al*., 2021). Shifts toward glutamate for both affinity and cooperativity estimates resulted in a reversal of the direction of probe dependence between species for MPEP, VU0424465 and DPFE, and introduced probe dependence towards glutamate for VU0409106, VU0409551 and VU0357121. As mentioned earlier, 50-80% of mGlu_5_ receptors are located on intracellular membrane in native cell preparations and subcellular localisation of mGlu_5_ can dictate subsequent receptor responses (Jong *et al*., 2009, 2019; Kumar *et al*., 2012; Vincent *et al*., 2016). DHPG is a membrane impermeable agonist and only activates cell surface receptors, therefore allosteric modulators are only acting on a discrete, membrane expressed pool of mGlu_5_ when modulating DHPG responses (Jong *et al*., 2005). Moreover, mGlu_5_ PAMs induce and modulate desensitisation and receptor internalisation of rat mGlu_5_, which could change the relative location of mGlu_5_ upon modulation (Hellyer *et al*., 2019; Arsova *et al*., 2021). How mGlu_5_ modulators affect receptor regulatory processes at the human receptor remains unknown. While overall expression of rat and human mGlu_5_ were similar in the current study, subcellular localisation and modulator effects on receptor regulatory processes were not assessed, with differences potentially playing a role in the different pharmacology evident between rat and human receptor.

Kinetic aspects of GPCR signalling and their relationship to ligand pharmacology are emerging as important considerations in the drug development pipeline (Hoare *et al*., 2018; Hoare, Tewson, Quinn, and Hughes, 2020). By understanding the effects of allosteric modulators on agonist kinetics, a deeper insight into mechanisms behind phenomena such as probe dependence and bias may be uncovered. While much work has been done on the effects of allosteric modulators on the binding kinetics of agonists at GPCRs, effects on agonist-induced signalling kinetics at a whole cell level remain understudied. Here we show mGlu_5_ PAMS have ligand dependent effects on the kinetics of glutamate and DHPG-induced iCa^2+^ mobilisation. The majority of PAMs, regardless of chemotype, increased initial rate of signalling for both agonists. Initial rate is derived from models used to describe enzyme kinetics and measures the rate of response prior to equilibrium, measured using the slope of the initial straight-line portion of an activation curve (Hoare *et al*., 2018; Hoare, Tewson, Quinn, Hughes, *et al*., 2020). Recent work on the M_2_ muscarinic receptor revealed a PAM enhanced initial rate of G protein association, reflective of stabilisation of the active state ternary complex (Burger *et al*., 2024). Conformational analysis of mGlu_5_ has also revealed PAMs stabilise conformations which facilitate G protein binding and active state formation (Nasrallah *et al*., 2021; Cannone *et al*., 2023; Li *et al*., 2024). Additionally, mGlu_2_ PAMs suppress sub-millisecond dynamic conformational changes in, lowering the energy barrier for glutamate to activate mGlu_2_ (Cao *et al*., 2021) and mGlu_5_ PAMs may enhance mGlu_5_ agonist-induced signalling rate through similar mechanisms, although further studies are required.

Interestingly, DPFE did not enhance initial rate of either agonist, and slowed the rate constant for the rise phase, indicating the potential for differential PAM effects based on chemotype. Perhaps the most striking differences in the current study were for the second site PAM, VU0357121, which did not affect glutamate kinetics but significantly increased DHPG initial rate. By binding to an as-yet unknown allosteric binding site, VU0357121 may stabilise a conformation preferentially activated by DHPG (Hammond *et al*., 2010). It should be noted VU0357121 displayed probe dependence towards glutamate, indicating a complicated relationship between kinetic analyses and probe dependence. However, it is clear mGlu_5_ PAMs differentially affected glutamate and DHPG signalling kinetics which we have previously linked to stabilisation of divergent activation states by glutamate and DHPG that differentially influence allosteric ligand-receptor interactions (Hellyer *et al*., 2020). Notably, some PAMs also affected the kinetics of the decay phase for both agonists. The decay phase represents a composite of multiple mechanisms related to signal cessation, including second messenger decay, iCa^2+^ clearance and receptor desensitisation (Hoare, Tewson, Quinn, Hughes, *et al*., 2020). mGlu_5_ PAMs, including those tested here, rapidly desensitise mGlu_5_ alone, and enhance agonist dependent mGlu_5_ desensitisation, which may be reflected in changes to iCa^2+^ signal decay (Hellyer *et al*., 2019). However, some PAMs slowed agonist signal decay and resulted in increased response duration and associated AUC, demonstrating the intricate interplay between receptor activation and regulatory mechanisms when considering kinetic of functional response.

Although NALs have mostly been used as tool compounds, mGlu_5_ NALs are gaining interest as potential treatments for Alzheimer’s disease (AD). ALX-001 (formerly BMS-984923) is NAL for mGlu_5_ mediated calcium signalling but inhibits mGlu_5_ interactions with cellular prion protein (PrP^C^) bound to amyloid beta (Aβo) (Huang *et al*., 2016; Haas *et al*., 2017). Pathogenic PrP^C^-Aβo-mGlu_5_ interactions are involved in AD pathogenesis, and ALX-001 disruption of this complex restores memory deficits in mouse models of AD, with a clinical trial (NCT04805983) now recruiting to move this compound forward into humans (Haas *et al*., 2017). Compounds inhibiting pathogenic mGlu_5_ signalling without affecting glutamate activity are highly desirable, as inhibition of glutamatergic mGlu_5_ activity can induce psychotomimetic effects and further impair learning and memory (Campbell *et al*., 2004; Porter *et al*., 2005; Jacob *et al*., 2009; Rodriguez *et al*., 2010; Um *et al*., 2013; Abou Farha *et al*., 2014; Gould *et al*., 2016; Holter *et al*., 2021). Encouragingly, initial ALX-001 pharmacological characterisation was undertaken using human mGlu_5_, with no evidence of NAM activity in iCa^2+^ mobilisation assays (Huang *et al*., 2016). Similarly, here 5MPEP was also a NAL for iCa^2+^ mobilisation and lacked agonist activity in IP_1_ accumulation assays at human mGlu_5._ However, a lack of activity in iCa^2+^ mobilisation assays does not preclude effects on other signalling endpoints, as biased neutral mGlu_5_ ligands have been identified that are inactive for calcium signalling but agonists for IP_1_ accumulation (Hellyer *et al*., 2018). In addition, NALs at mGlu_5_ may have activity at other mGlu subtypes and Class C GPCRs, as known Class C ligands have NAL activity at mGlu_5_ (Hellyer *et al*., 2018). Unknown and potentially unappreciated modulatory or agonist activity for NALs across divergent pathways highlights caution must be used when progressing mGlu_5_ NALs through screening and preclinical testing, and even the use of the term “neutral” must be recognised as context-, species and assay-dependent. Rigorous pharmacological characterisation is crucial in novel and ongoing mGlu_5_ drug discovery programs as NALs gain traction as potential therapeutics.

In summary, we have determined the signalling fingerprints of eleven mGlu_5_ allosteric modulators in recombinant cells expressing human mGlu_5_. Our data contribute to an emerging body of evidence aimed at understanding the inherent complexity in quantifying allosteric modulator pharmacology, with emphasis on the importance of species-differences in ligand pharmacology. Here we show ligand-dependent species differences in the molecular pharmacological properties among mGlu_5_ allosteric modulators. Changes to functional affinity, cooperativity estimates and probe dependence were evident for select ligands. Understanding potential for differences in pharmacological parameters is important for predicting preclinical efficacy; for example PAM cooperativity rather than affinity was correlated with efficacy in rodent models of psychosis (Gregory *et al*., 2019). Additionally, mGlu_5_ NAM affinity is related to receptor occupancy and brain exposure-response relationships across species, including in humans (Bennett *et al*., 2021). Of the ligands with differential pharmacology, VU0424465 is a robust ago-PAM with seizurogenic properties unlikely to be tested in humans, and as such the differences seen here are unlikely to impact translatability (Rook *et al*., 2013). However, VU0409106 has gained interest as a potential treatment for binge-eating, displaying differences in both binding and functional affinity between species that could potentially impact its activity in patients (Oliveira *et al*., 2021). While the only clinically tested compound in the current study, fenobam, showed no species differences, it is important to test existing and novel clinical candidates at human mGlu_5_ to identify potential differences in pharmacology that may impact future translation of mGlu_5_ allosteric modulators into the clinic.

## CRediT Authorship Contributions

**Razzak:** conceptualization, data curation, investigation, formal analysis, writing – original draft, writing – reviewing and editing; **Tran:** investigation, formal analysis, writing – reviewing and editing; **McCague:** investigation, formal analysis, writing – reviewing and editing; **Sengmany:** investigation, formal analysis, writing – reviewing and editing; **Kos:** investigation, formal analysis, writing – reviewing and editing; **Langiu:** investigation, formal analysis, writing – reviewing and editing; **Li:** conceptualization, writing – reviewing and editing; **Hellyer:** investigation, formal analysis, writing – original draft, writing – reviewing and editing, conceptualization, supervision; **Gregory:** formal analysis, writing – original draft, writing – reviewing and editing, funding acquisition, conceptualization, supervision, project administration

## Abbreviations

7TM: 7 transmembrane domain
CNS: central nervous system
DMEM: Dulbecco’s modified Eagle’s medium
FBS: foetal bovine serum
GPCR: G protein-coupled receptor
HBSS: Hank’s Balanced Salt Solution
HEK293A: human embryonic kidney 293
iCa^2+^: intracellular calcium
IP_1_: inositol 1-phosphate
mGlu: metabotropic glutamate receptor
MPEP: 2-Methyl-6-(phenylethynyl)pyridine
NAL: neutral allosteric ligand
NAM: negative allosteric modulator
PAM: positive allosteric modulator
SAR: structure-activity relationship.

